# MerTK-dependent efferocytosis by monocytic-MDSCs mediates resolution of post-lung transplant injury

**DOI:** 10.1101/2024.01.18.576261

**Authors:** Victoria Leroy, Denny J. Manual Kollareth, Zhenxiao Tu, Jeff Arni C. Valisno, Makena Woolet-Stockton, Biplab Saha, Amir M. Emtiazjoo, Mindaugas Rackauskas, Lyle L. Moldawer, Philip A. Efron, Guoshuai Cai, Carl Atkinson, Gilbert R. Upchurch, Ashish K. Sharma

## Abstract

**ABSTRACT:** *Rationale:* Patients with end stage lung diseases require lung transplantation (LTx) that can be impeded by ischemia-reperfusion injury (IRI) leading to subsequent chronic lung allograft dysfunction (CLAD) and inadequate outcomes.

*Objectives:* We examined the undefined role of MerTK (receptor Mer tyrosine kinase) on monocytic myeloid-derived suppressor cells (M-MDSCs) in efferocytosis (phagocytosis of apoptotic cells) to facilitate resolution of lung IRI.

*Methods:* Single-cell RNA sequencing of lung tissue and BAL from post-LTx patients was analyzed. Murine lung hilar ligation and allogeneic orthotopic LTx models of IRI were used with Balb/c (WT), *cebpb*^-/-^ (MDSC-deficient), *Mertk^-/-^* or MerTK-CR (cleavage resistant) mice. Lung function, IRI (inflammatory cytokine and myeloperoxidase expression, immunohistology for neutrophil infiltration), and flow cytometry of lung tissue for efferocytosis of apoptotic neutrophils were assessed in mice.

*Measurements and Main Results:* A significant downregulation in MerTK-related efferocytosis genes in M-MDSC populations of CLAD patients compared to healthy subjects was observed. In the murine IRI model, significant increase in M-MDSCs, MerTK expression and efferocytosis was observed in WT mice during resolution phase that was absent in *cebpb^-/-^* Land *Mertk^-/-^* mice. Adoptive transfer of M-MDSCs in *cebpb^-/-^* mice significantly attenuated lung dysfunction, and inflammation leading to resolution of IRI. Additionally, in a preclinical murine orthotopic LTx model, increases in M-MDSCs were associated with resolution of lung IRI in the transplant recipients. *In vitro* studies demonstrated the ability of M-MDSCs to efferocytose apoptotic neutrophils in a MerTK-dependent manner.

*Conclusions:* Our results suggest that MerTK-dependent efferocytosis by M-MDSCs can significantly contribute to the resolution of post-LTx IRI.

## INTRODUCTION

For patients with advanced end stage lung diseases, the ultimate option is lung transplantation (LTx) (1). However, post-LTx ischemia-reperfusion injury (LTx-IRI) can lead to primary graft dysfunction (PGD) and chronic rejection (CLAD) without effective therapeutic modalities (1-3). IRI is characterized by immune cell infiltration and activation, increased vascular permeability, and production of inflammatory mediators, including reactive oxygen species (4). Dysregulation of endogenous mechanisms of inflammation-resolution lead to exacerbated tissue injury and graft dysfunction (5, 6). Since PGD development is a primary risk factor for CLAD, it is imperative, to understand endogenous mechanisms of inflammation-resolution that can be harnessed to facilitate graft acceptance and tissue homeostasis.

Broadly, the inflammation-resolution process is hallmarked by a variety of factors that include the presence and activation of immunosuppressive cells, the cessation of pro-inflammatory cell infiltration, and subsequent clearance of apoptotic cells (efferocytosis) to prevent secondary necrosis (7). Among immunosuppressive cell populations, myeloid-derived suppressor cells (MDSCs) have garnered research interest in the transplantation field both has a therapeutic measure as well as a diagnostic biomarker (8-10). This heterogenous cell population, comprising of monocytic-(M-) and granulocytic (G- or PMN-)- MDSCs are notable for their myriad of immunosuppressive/pro-resolving actions including modulation of cytokines, exhaustion of pro-inflammatory cells, and promotion of pro-resolving cell phenotypes (11-14). A pivotal characteristic of the resolution phase is efferocytosis, which involves the clearance of apoptotic cells and debris (15). This key pro-resolving action is carried out by both professional and non-professional phagocytes at the direction of various pro-efferocytic receptors including Mer proto-oncogene tyrosine kinase (MerTK) (16, 17). When efferocytosis is dysregulated, apoptotic cells remain unengulfed and quickly become necrotic and exacerbate the inflammatory response (18). However, the role of efferocytosis and dysregulation of inflammation-resolution pathways via MerTK in post-LTx IRI remains to be delineated.

In this study, we investigate if M-MDSCs contribute to the resolution of lung IRI via MerTK-dependent efferocytosis leading to graft survival. These results signify the importance of dysregulation of inflammation-resolution due to MerTK-cleavage and subsequent efferocytosis that contributes to exacerbated tissue inflammation and primary graft dysfunction.

## METHODS

Details available in the online data supplement.

### Human BAL analysis

Patients undergoing lung transplantation at University of Florida Health were consented for bronchoalveolar lavage (BAL) fluid collected in accordance with the University of Florida Institutional Review Board (#IRB201900987).

### Human single cell RNA sequencing analysis

Single-cell RNA sequencing data, previously described, was analyzed for myeloid-cell populations and subsequent differential gene expression (19).

### Lung IR injury model

Lung IRI via hilar ligation was performed in male Balb/c (wild-type; WT), *cebpb^-/-^,* C57Bl/6, *Mertk^-/-^,* and MerTK-Cleavage Resistant (MerTK-CR) mice, as we have previously described (20, 21).

### Brain dead (BD) orthotopic lung transplantation

Murine LTx was performed on brain dead (BD) donors using C57Bl/6 donor and Balb/c recipient mice for 24- or 72hrs of reperfusion, as previously described (22-24).

### Statistical Analysis

Values are presented as the mean ± standard error of the mean (SEM) and statistical evaluation was performed with GraphPad Prism 10 software (GraphPad, La Jolla, CA).

## RESULTS

### Efferocytosis-related genes are downregulated in M-MDSCs in CLAD patients

We analyzed single-cell RNA sequencing data from CLAD patients (*n*=4) and donor tissue (DT; *n*=3) from a recently reported study to identify differences in M-MDSC cell populations from myeloid cell-specific clusters (19). After data normalization and principal component analysis, we identified a total of 18 myeloid cell clusters using uniform manifold approximation and projection (UMAP) (**Figure 1A**). Subcluster analysis of these myeloid populations revealed Cluster 4 as M-MDSCs based on expression of *HLA-DRA, ITGAM, CD33, CD14, FUT4, IL-10, VEGFA,* as previously described (**Figure E1**) (25, 26). M-MDSCs, which display a variety of immunosuppressive capabilities, and therefore should contribute to graft tolerance were present in patients with chronic rejection, which prompted further examination.

**Figure 1.**
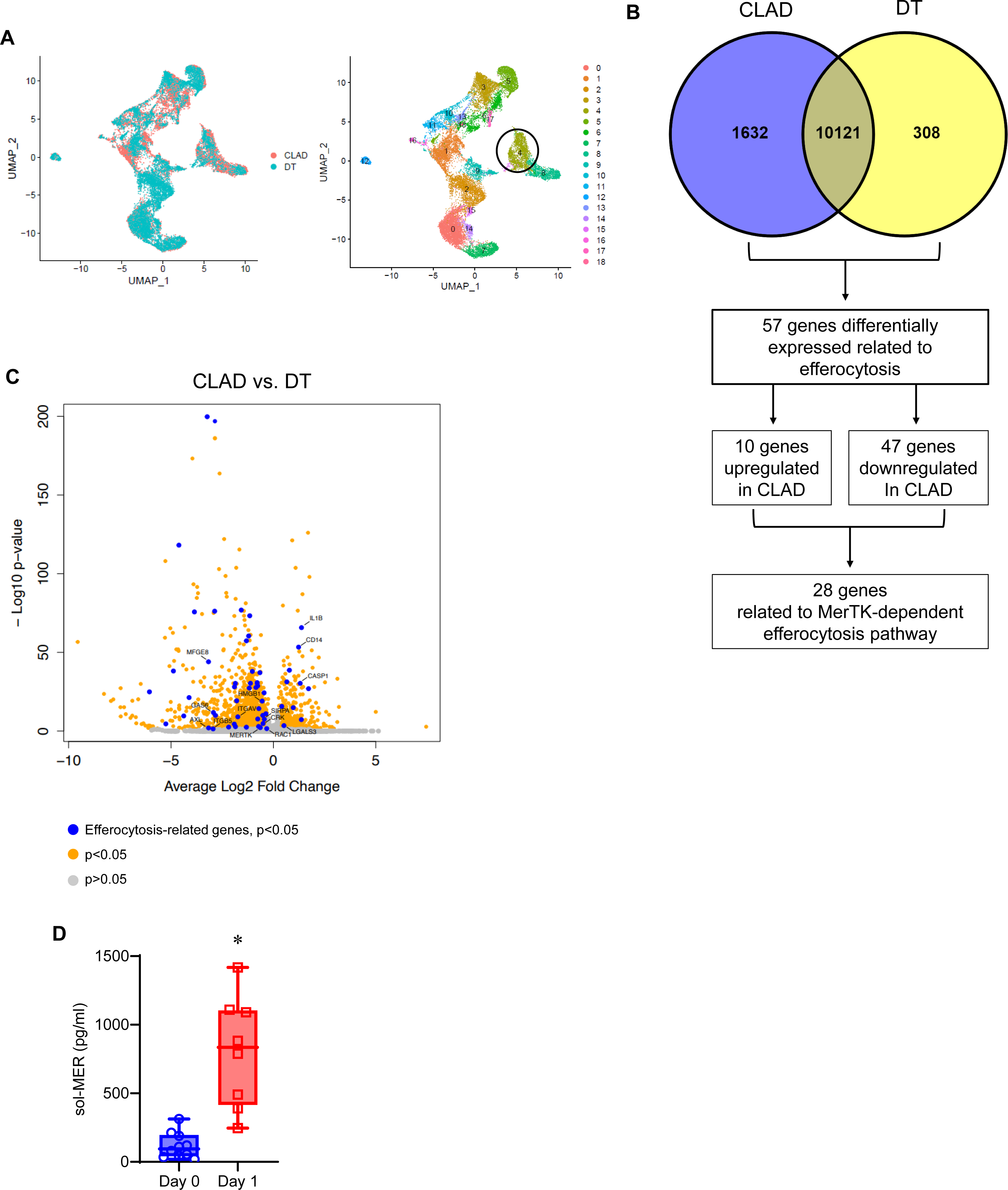
Single-cell RNA-sequencing analysis of myeloid cells reveals downregulation of efferocytosis related genes in M-MDSCs of CLAD patients. **(A)** UMAP visualization of 18 myeloid cell clusters in lung tissue of CLAD patients and donor controls. **(B)** Venn diagram outlining identification of differentially expressed genes (DEGs) for MerTK-dependent efferocytosis. Downregulated genes in CLAD (blue) and donor tissue (DT; yellow) are described. **(C)** Volcano plot of DEGs of myeloid cell cluster 4. Genes identified by blue dots are efferocytosis-related genes with differential expression of p<0.05. Orange dots are other DEGs with p<0.05. Grey dots are genes that are not significant (p>0.05). **(D)** Quantification of sol-MER levels in BAL of patients showed significant increase on day 1 post-LTx compared to day 0. *p<0.0001; n=8-10/group.

Analysis of differentially expressed genes (DEGs) in M-MDSCs detected 1,632 genes downregulated in CLAD patients and 308 genes downregulated in DT. In depth analysis revealed 5 of the top 30 DEGs (*CD163, C25H, PELI1, IL10, VSIG4)* were related to a downregulation in efferocytosis (surveyed from genes outlined in **Table E1**). The efferocytosis process, or clearance of apoptotic cells, is vital to inflammation-resolution and especially crucial in pulmonary IRI (27). Since PGD can contribute to CLAD, we further explored efferocytosis-related genes in patients with CLAD. Of the 57 ERGs, 47 were downregulated in CLAD patients, the majority of which were directly related to MER proto-oncogene, tyrosine kinase (MerTK)-mediated efferocytosis (**Figure 1B-C; Table E2**), which includes *MERTK, AXL, GAS6, ADAM9, SIRP*α*, CASP1*, and *RAC1* among others (**Table E3**). MerTK, as a cell surface receptor is subject to cleavage, which contributes to defective efferocytosis (28). The cleavage of MerTK into soluble MER (sol-MER) can be quantified in patient BAL. Therefore, we also analyzed patient BAL on days 0 and 1 post-LTx to gain insight into sol-MER levels at the critical period in which PGD typically arises. Compared to day 0 controls, sol-MER was significantly increased in patients on day 1 post-LTx (802.0±142.7 vs. 117.5±29.8 pg/mL; p<0.0001; **Figure 1D**). These clinical findings prompted us to further explore the endogenous mechanisms associated with MerTK-dependent efferocytosis, via M-MDSCs, in our preclinical models of lung IRI.

### M-MDSCs mediate the endogenous resolution of experimental lung IRI

Using the murine hilar ligation model of lung IRI (**Figure 2A**), we observed that maximal lung dysfunction occurred at IRI (6hrs) compared to sham controls as seen by decreased pulmonary compliance (2.9±0.1 vs. 6.2±0.2 µl/cm H_2_O; p<0.0001), increased airway resistance (1.5±0.1 vs. 0.6±0.04 cm H_2_O/µl/sec; p<0.0001) and pulmonary artery (PA) pressure (12.7±0.4 vs. 5.6±0.1 cm H_2_O; p<0.0001). Furthermore, lung dysfunction was attenuated at IRI (24hrs) compared to IRI (6hrs) indicating endogenous resolution of lung dysfunction (**Figure 2B-D**). Next, we investigated the contribution of M-MDSCs to inflammation resolution and lung injury attenuation using *cebpb^-/-^* (MDSC knockout) mice. Following IRI (6hrs), there was substantial lung dysfunction in *cebpb^-/-^*compared to respective sham control as seen by significant decrease in pulmonary compliance (2.8±0.3 vs. 6.5±0.2 µl/cm H_2_O; p<0.0001) and significant increase in airway resistance (1.3±0.2 vs. 0.61±0.04 cm H_2_O/µl/sec; p<0.0013) and pulmonary artery (PA) pressure (11.8±0.6 vs. 5.7±0.2 cm H_2_O; p<0.0001). Contrary to WT mice, sustained lung dysfunction was observed in *cebpb^-/-^* mice following IRI (24hrs) with significantly decreased pulmonary compliance (3.1±0.1 vs. 5.2±0.3 µl/cm H_2_O; p<0.0001) and increased airway resistance (1.2±0.2 vs. 0.8±0.05 cm H_2_O/µl/sec; p<0.05) and PA pressure (11.2±0.6 vs. 7.1±0.2 cm H_2_O; p<0.0001) compared to WT mice indicating dysregulated resolution (**Figure 2B-D**). There was no difference in lung function among all WT and *cebpb^-/-^* sham groups (data not shown).

**Figure 2.**
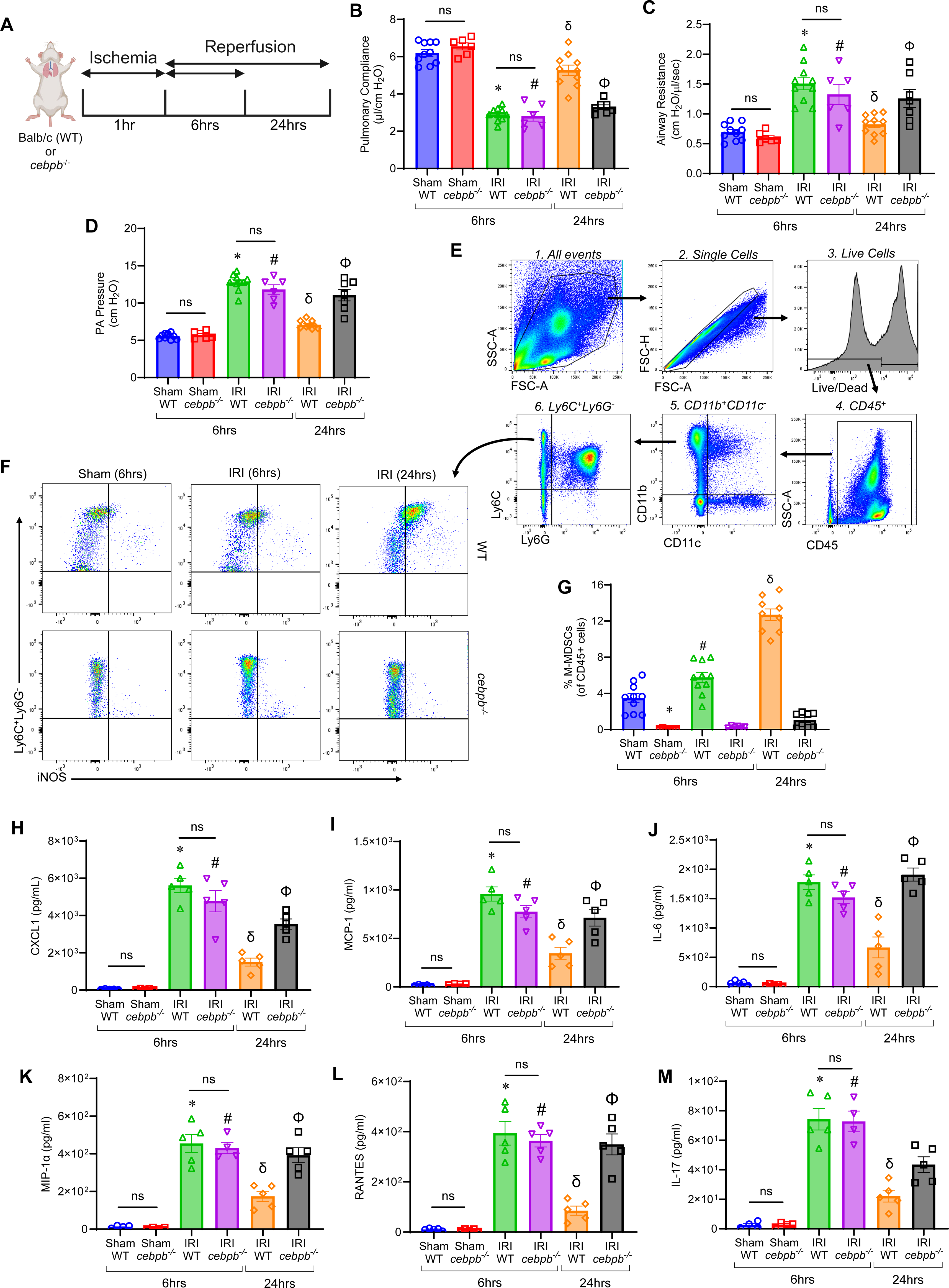
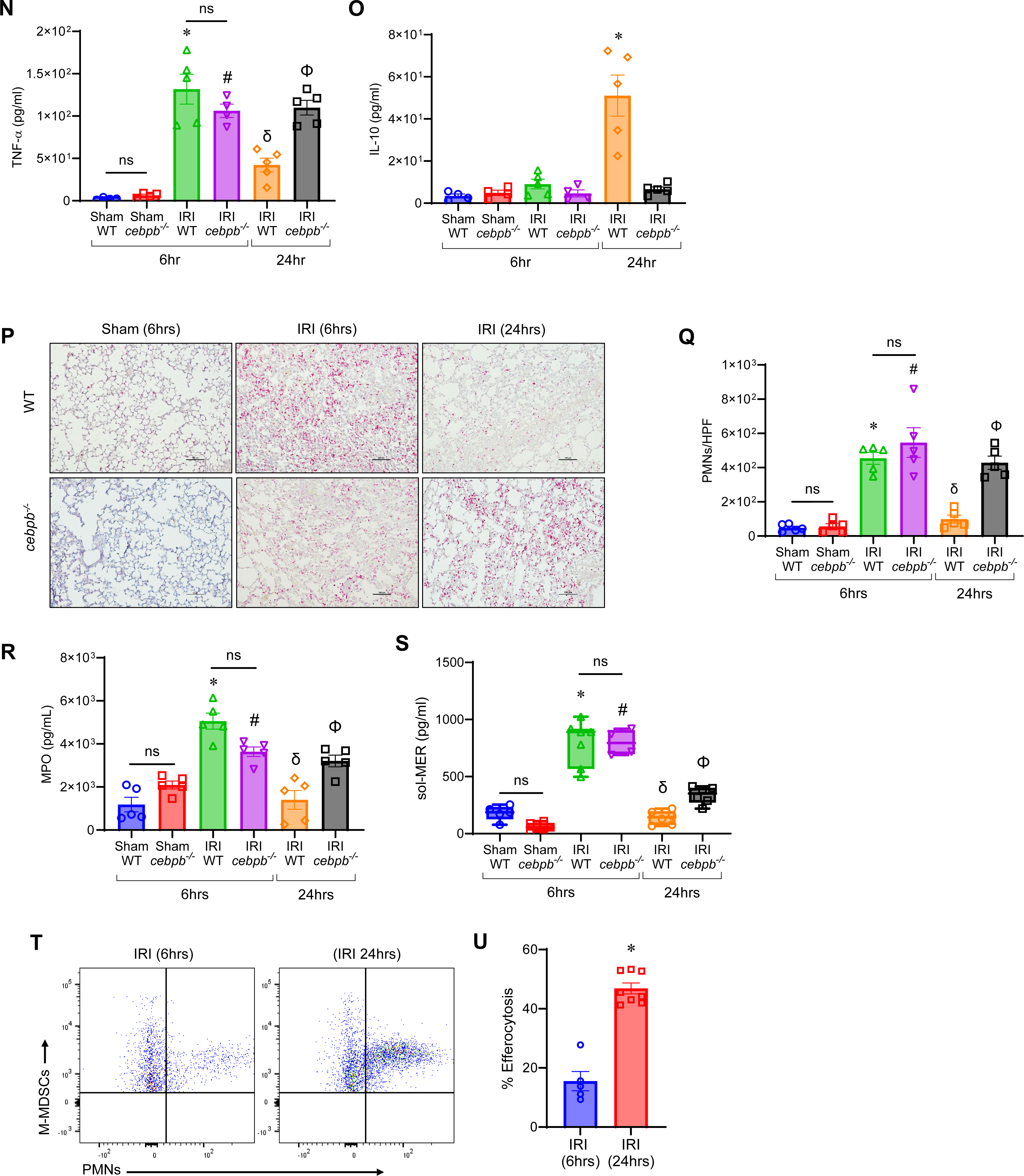
Increase in M-MDSCs is associated with the endogenous resolution of lung IRI. **(A)** Representative schematic depicting the hilar ligation IRI model where left lung undergoes 1hr of ischemia followed by 6- or 24hrs-of reperfusion in Balb/c (WT) and *cebpb^-/-^* mice. **(B-D)** Significant lung dysfunction was observed in WT and *cebpb^-/-^* mice following 6hrs compared to sham controls that was mitigated after 24hrs. However, lung dysfunction was significantly higher in *cebpb^-/-^* mice compared to WT mice after 24hrs. ns, not significant; *p<0.0001 vs. WT sham; ^#^p<0.0001 vs *cebpb^-/-^* sham; ^δ^p<0.0001 vs. WT IRI (6hrs); ^Φ^p<0.001 vs. WT IRI (24hrs); *n*=6-10/group. **(E-G)** Representative flow cytometry gating strategy for M-MDSC identification in lung tissue. The percentage of M-MDSCs was significantly upregulated in WT mice following IRI (24hrs) compared to IRI (6hrs) or sham mice. A markedly significant mitigation of M-MDSCs was observed in *cebpb^-/-^* compared to WT mice. *p=0.005 vs. WT sham; ^#^p<0.009 vs WT sham and *cebpb^-/-^* IRI (6hrs); ^δ^p<0.0001 vs. WT IRI (6hrs) and *cebpb^-/-^* IRI (24hrs); *n*=6-10/group. **(H-O)** Pro-inflammatory cytokine and chemokine levels were significantly elevated in WT and *cebpb^-/-^* mice after 6hrs of IRI compared to sham controls, and were mitigated in WT mice following 24hrs, but not in *cebpb^-/-^* mice. Anti-inflammatory IL-10 expression was significantly increased in WT mice following 24hrs compared to all other groups. ns, not significant; *p<0.05 vs. WT sham; ^#^p<0.05 vs *cebpb^-/-^* sham; ^δ^p<0.05 vs. WT IRI (6hrs); ^Φ^p<0.05 vs. WT IRI (24hrs); *n*=5/group. **(P-R)** PMN infiltration and activation were significantly increased in WT and *cebpb^-/-^* mice after 6hrs compared to sham controls, which was attenuated in WT mice following 24hrs but not in *cebpb^-/-^*mice. ns, not significant; *p<0.05 vs. WT sham; ^#^p<0.05 vs *cebpb^-/-^* sham; ^δ^p<0.05 vs. WT IRI (6hrs); ^Φ^p<0.05 vs. WT IRI (24hrs); *n*=5/group. PMNs were quantified per high power field (HPF). Scale bars represent 100µm. **(S)** sol-MER levels were significantly increased in both WT and *cebpb^-/-^* after 6hrs compared to respective sham controls. These levels were mitigated in WT mice after 24hrs, but not in *cebpb^-/-^*mice. ns, not significant; *p<0.0007 vs. WT sham; ^#^p<0.0001 vs *cebpb^-/-^* sham; ^δ^p<0.0001 vs. WT IRI (6hrs); ^Φ^p<0.05 vs. WT IRI (24hrs); *n*=4-7/group. **(T-U)** Efferocytosis of neutrophils by M-MDSCs, measured via flow cytometry, was significantly increased after 24hrs compared to 6hrs in WT mice following IRI. *p=0.0016 vs. IRI (6hrs); *n*=5-8/group.

Interestingly, WT mice displayed a significant increase in the M-MDSC population (CD45^+^CD11b^+^CD11c^-^Ly6G^-^Ly6C^+^iNOS^+^) infiltrating in the lungs during resolution phase (24hrs) compared to inflammation phase (6hrs) post-IRI (12.7±0.6 vs. 5.7±0.5%; p<0.0001; **Figure 2E-G**). There was a significant decrease in M-MDSCs in *cebpb^-/-^* mice compared to WT mice after IRI (24hrs) (1.0±0.3 vs. 12.7±0.6%; p<0.0001; **Figure 2E- G**). These results highlight a critical association between M-MDSC infiltration and inflammation-resolution.

The dysfunction seen after IRI (6hrs) in WT and *cebpb^-/-^* mice was accompanied by an increase in proinflammatory cytokines (CXCL1, MCP-1, IL-6, MIP-1α, RANTES, IL-17, and TNF-α) as well as a decrease in anti-inflammatory IL-10 secretion in BAL (**Figure 2H-O**). Polymorphonuclear neutrophil (PMN) infiltration and activation (measured by myeloperoxidase; MPO) was elevated in both WT and *cebpb^-/-^* at IRI (6hrs) compared to respective sham controls (**Figure 2P-R**). PMN infiltration was significantly increased in *cebpb^-/-^* mice compared to WT mice at 24hrs-IRI (428.4±38.5 vs. 99.0±23.1 PMNs/HPF; p=0.0002) as well as activation (3207±269.6 vs. 1400±438.5 pg/mL; p=0.005) indicating dysregulation of efferocytosis in *cebpb^-/-^* mice.

To further explain the differences seen in PMN infiltration and activation, we investigated the processes that regulate the endogenous clearance of these cells. Both WT and *cebpb^-/-^* demonstrated increased sol-MER levels compared to respective sham controls after IRI. sol-MER levels were significantly decreased in WT mice following IRI (24hrs) compared to IRI (6hrs) (139.9±24.6 vs. 796.2±73.5 pg/mL; p<0.0001; **Figure 2S**). However, *cebpb^-/-^* mice displayed elevated sol-MER levels compared to WT at 24hrs (337.3±31.0 vs. 139.9±24.6 pg/mL; p=0.04; **Figure 2S**). mRNA expression of *MerTK* in lung tissue was significantly increased following 24hrs in WT mice compared to 6hrs (*see* **Figure E2** in the online supplement). Importantly, the endogenous efferocytosis by M-MDSCs in WT mice was significantly upregulated following IRI (24hrs) compared to IRI (6hrs) (46.9±1.9 vs. 15.5±3.2 %; p=0.0016; **Figure 2T-U**). Taken together, these data suggest that M-MDSCs are critical to the inherent resolution of lung IRI and mitigate inflammatory cytokine secretion, leukocyte trafficking, and efferocytosis.

### Adoptive transfer of M-MDSCs attenuates pulmonary dysfunction after lung IRI

To elucidate the reparative role of M-MDSCs in inflammation-resolution, we performed adoptive transfer of exogenous M-MDSCs (generated *in vitro*; **Figure E3**) (29) in WT mice prior to IRI (6hrs) (**Figure 3A**). WT mice treated with M-MDSCs showed significant improvement in lung function compared to untreated mice after IRI (6hrs) as demonstrated by an increase in pulmonary compliance (4.1±0.2 vs. 2.5±0.2 µl/cm H_2_O; p<0.001) as well as decrease in airway resistance (1.0±0.04 vs. 1.4±0.1 cm H_2_O/µl/sec; p<0.001) and PA pressure (8.8±0.5 vs. 12.7±0.4 cm H_2_O; p<0.003) (**Figure 3B-D**).

**Figure 3.**
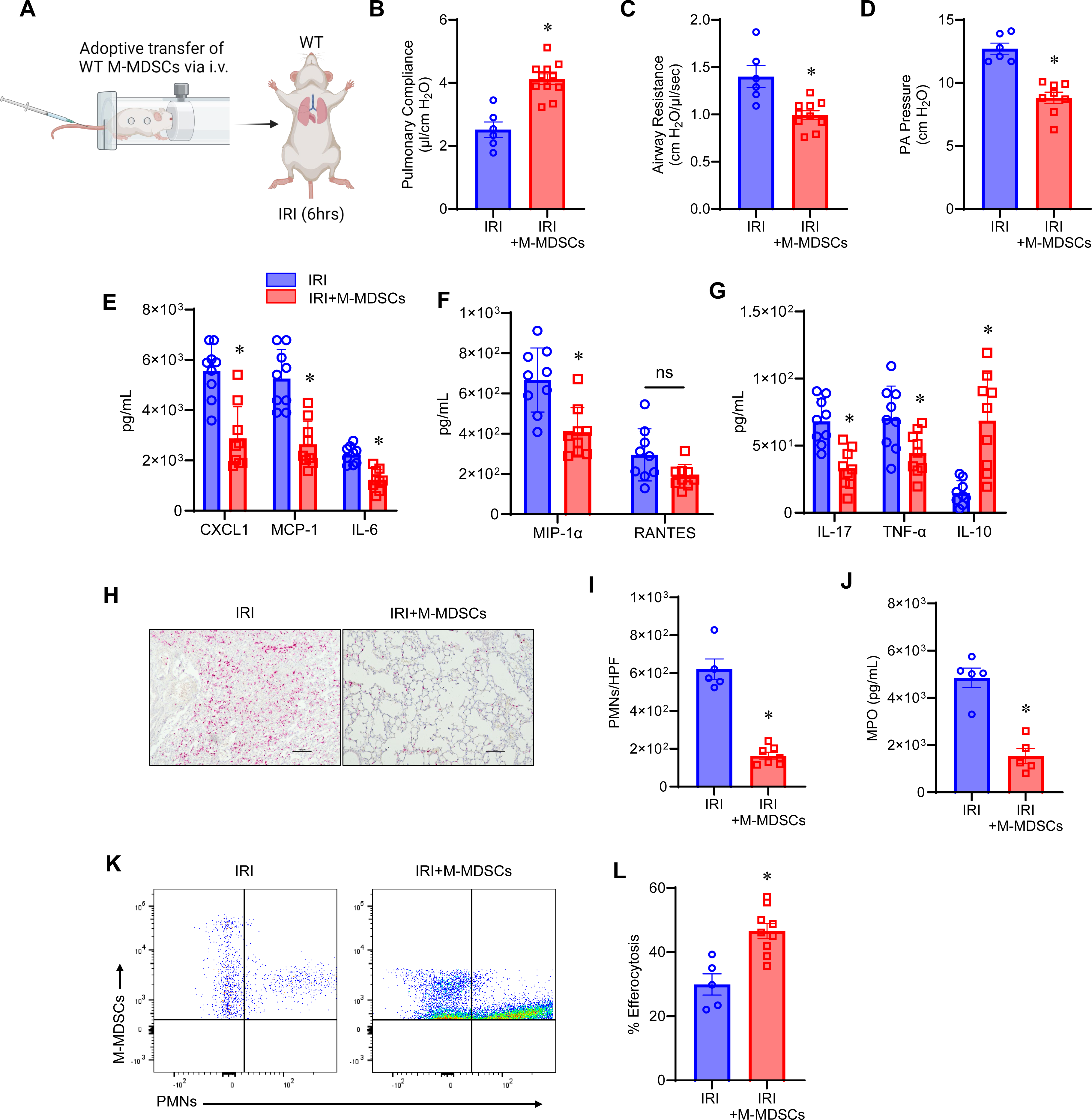
Administration of M-MDSCs attenuates pulmonary dysfunction after IRI. **(A)** Schematic depicting adoptive transfer of WT M-MDSCs prior to IRI (6hrs) in WT mice. **(B-D)** Treatment with M-MDSCs significantly improved lung dysfunction compared to untreated mice. *p<0.003 vs. IRI alone; *n*=6-11/group. **(E-G)** Pro-inflammatory cytokines were significantly reduced and anti-inflammatory IL-10 expression was significantly increased in M-MDSC-treated mice compared to untreated mice. ns, not significant *p<0.05 vs. IRI alone; *n*=9/group. **(H)** Representative images of histological staining for PMNs after IRI with and without M-MDSC treatment. Scale bars represent 100µm. **(I-J)** Neutrophil infiltration and MPO levels were significantly attenuated in mice treated with M-MDSCs. *p<0.008 vs. IRI alone; *n*=5/group. **(K-L)** Adoptively transferred M-MDSCs demonstrated a significant increase in efferocytosis of apoptotic neutrophils. *p=0.004 vs. IRI alone; *n*=5-9/group.

Additionally, M-MDSC treated mice had significantly less proinflammatory cytokines and increased anti-inflammatory/pro-resolving cytokine, IL-10, compared to untreated mice after IRI (6hrs) (**Figure 3E-G**). Moreover, M-MDSC treated mice had significantly decreased neutrophil infiltration (163.7±18.2 vs. 620.0±54.3 PMNs/HPF; p=0.0025) **(Figure 3H-I**) and MPO levels (1533.0±318.0 vs. 4851.0±410.0 pg/mL; p=0.0079) (**Figure 3J**) compared to untreated controls, suggesting increased efferocytosis of endogenous neutrophils by exogenously administered M-MDSCs. Adoptively transferred M-MDSCs also significantly increased efferocytosis of exogenously administered apoptotic PMNs compared to endogenous M-MDSCs after IRI (6hrs) (46.5±2.4 vs. 29.9±3.2 %; p=0.004) (**Figure 3K-L**). These findings suggest that exogenously administered M-MDSCs are capable of mitigating early pulmonary IRI.

### MerTK is critical to the endogenous resolution of lung IRI

To understand the role of MerTK during efferocytosis and inflammation-resolution, we analyzed *Mertk^-/-^* mice using the lung IRI model. *Mertk^-/-^* mice subject to IRI (24hrs) failed to resolve with a significant increase in lung dysfunction compared to C57Bl/6 mice as seen by increased pulmonary compliance (2.7±1.7 vs. 5.4±0.2 µl/cm H_2_O; p<0.0001), as well as increased airway resistance (1.5±0.1 vs. 0.7±0.1 cm H_2_O/µl/sec; p<0.0001) and PA pressure (10.7±0.4 vs. 6.3±0.3 cm H_2_O; p<0.0001; **Figure E4**). This lung dysfunction was accompanied by a significant increase in proinflammatory cytokine and chemokine expression and significant decrease in IL-10 in BAL fluid from these mice compared to C57Bl/6 mice (24hrs) (**Figure E5**). Moreover, the *Mertk^-/-^* mice displayed significantly increased PMN infiltration compared to C57Bl/6 after IRI (24hrs) (506.8±30.9 vs. 54.8±17.4 PMNs/HPF; p<0.0001; **Figure E6A-B**) and MPO expression (5321±469.2 vs. 2320±668.9 pg/mL; p=0.01; **Figure E6C**), indicating dysregulated efferocytosis of PMNs in these loss of function mice.

Additionally, we sought to examine the role of MerTK by assessing injury resolution in MerTK-cleavage resistant (MerTK-CR) mice (30). MerTK-CR mice were significantly protected after IRI (6hrs) compared to C57Bl/6 counterparts by increased pulmonary compliance (3.6±0.2 vs. 2.3±0.2 µl/cm H_2_O; p=0.007) as well as decreased airway resistance (1.0±0.1 vs. 1.7±0.03 cm H_2_O/µl/sec; p<0.0001) and PA pressure (8.8±0.4 vs. 13.6±0.4 cm H_2_O; p<0.0001; **Figure E7**). MerTK-CR mice demonstrated significant protection via a decrease in proinflammatory cytokines and chemokines and increase in anti-inflammatory IL-10 compared to IRI (6hrs) in C57Bl/6 mice (**Figure E8**). MerTK-CR mice displayed enhanced efferocytosis after IRI (6hrs) compared to C57Bl/6 which was evidenced by a significant decrease in PMN infiltration (148.9±13.14 vs. 680.4±44.24 PMNs/HPF; p<0.0001; **Figure E9A-B**) and MPO expression (1962±291.5 vs. 4591±660.2 pg/mL; p=0.0006; **Figure E9C**). The expression of MerTK was absent in *Mertk^-/-^* mice and detectable in MerTK-CR mice (**Figure E10A-B**). Taken together, these results demonstrate the critical role of MerTK in the efferocytosis-mediated resolution of lung IRI.

### MerTK-mediated efferocytosis by M-MDSCs contributes to inflammation-resolution during lung IRI

Next, we sought to understand the involvement of MerTK on M-MDSCs in the resolution of lung IRI. We performed adoptive transfer of WT and *Mertk^-/-^* M-MDSCs into *cebpb^-/-^* prior to IRI (24hrs) (**Figure 4A**). *cebpb^-/-^* mice treated with WT M-MDSCs demonstrated significant protection in lung dysfunction following IRI (24hrs) that was absent in untreated counterparts as observed by increased pulmonary compliance (3.8±0.3 vs. 2.5±0.1 µl/cm H_2_O) as well as decreased airway resistance (1.1±0.1 vs. 1.6±0.1 cm H_2_O/µl/sec) and PA pressure (9.0±0.5 vs. 12.2±0.3 cm H_2_O). However, *cebpb*^-/-^ mice treated with *Mertk^-/-^* M-MDSCs showed significantly increased lung dysfunction and injury compared to mice treated with WT M-MDSCs which were equivalent to untreated counterparts **(Figure 4B-D**). Additionally, pro-inflammatory cytokines and chemokines were significantly mitigated in mice treated with WT M-MDSCs, but not with *Mertk^-/-^*M-MDSCs (**Figure 4E-G**). PMN infiltration following IRI (24hrs) was mitigated in mice treated with WT M-MDSCs compared to *Mertk^-/-^* M-MDSC treated mice (163.7±18.2 vs. 409.4±33.1 PMNs/HPF PMNs/HPF; p=0.0005), as well as MPO expression (2380.0±393.7 vs.3873±273.2 pg/mL; p=0.04) (**Figure 4J**).

**Figure 4.**
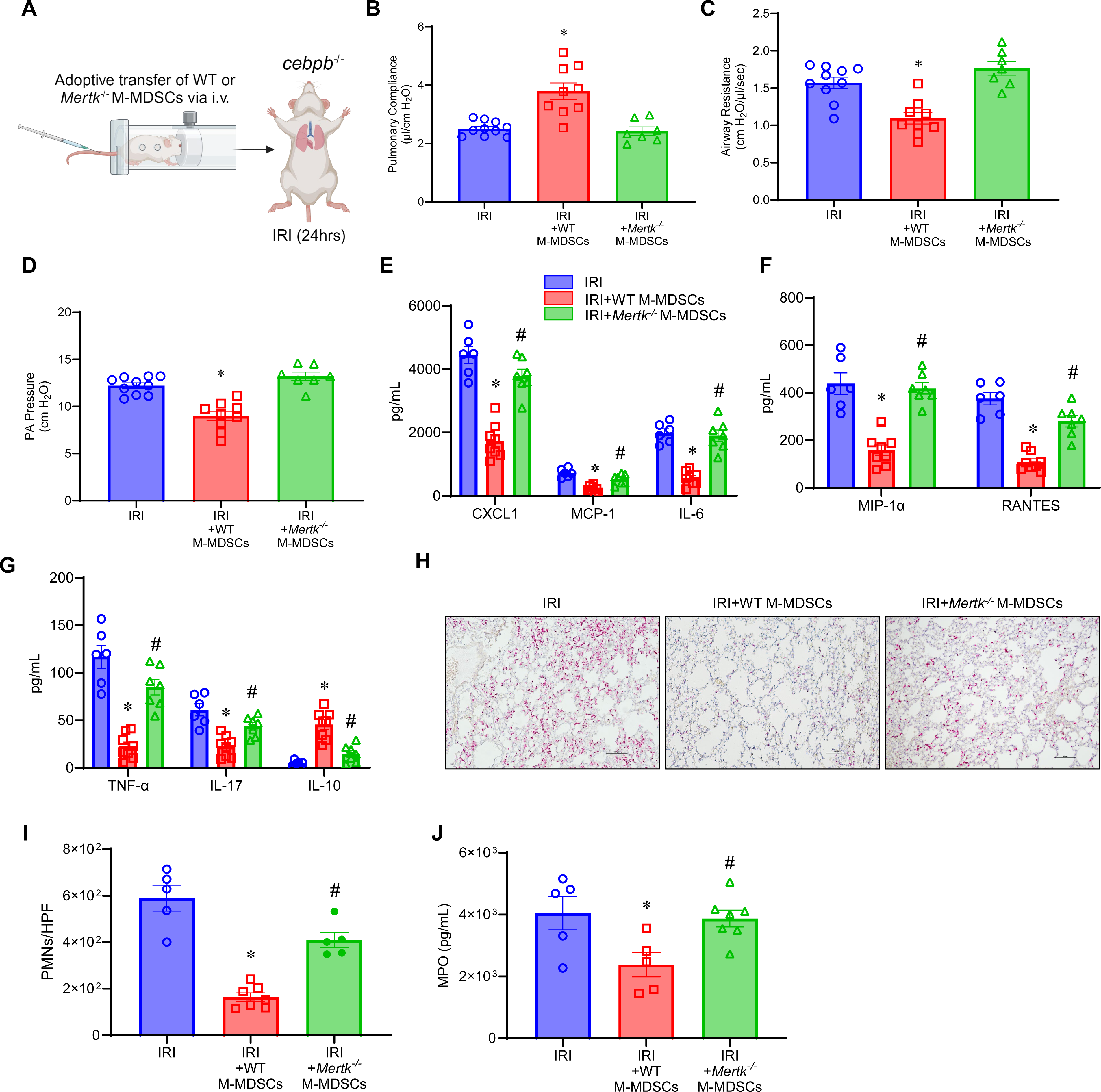
M-MDSC mediated resolution of lung IRI is mediated via MerTK. **(A)** Schematic depicting adoptive transfer of M-MDSCs prior to IRI (24hrs) in *cebpb^-/-^* mice. **(B-D)** Treatment with M-MDSCs from WT mice, but not from *Mertk^-/-^*, significantly mitigated lung dysfunction compared to untreated mice. ns, not significant; *p<0.006 vs. IRI all other groups; *n*=7-9/group. **(E-G)** Expression of pro inflammatory cytokines were significantly decreased in mice treated with WT M-MDSCs, but not with *Mertk^-/-^* M- MDSCs. *p<0.05 vs. IRI alone; ^#^p<0.05 vs WT M-MDSC treated mice; *n*=6-8/group. **(H-I)** PMN infiltration was significantly attenuated by treatment with WT-derived M-MDSCs but not *Mertk^-/-^*-derived M-MDSCs. Scale bars represent 100µm. *p<0.0001 vs. IRI alone; ^#^p=0.0005 vs WT M-MDSC treated mice; *n*=5-7/group. **(J)** MPO levels in BAL were significantly mitigated by WT M-MDSCs, but not by *Mertk^-/-^* M-MDSCs. *p<0.05 vs. IRI alone; ^#^p<0.05 vs WT M-MDSC treated mice; *n*=5-7/group.

### Apoptotic PMNs undergo efferocytosis by M-MDSCs in a MerTK-dependent manner

The integral role of MerTK-dependent efferocytosis by M-MDSCs was further investigated by our *in vitro* studies (**Figure 5A**). Confocal analysis of M-MDSCs co-cultured with apoptotic PMNs demonstrated co-expression of ingested PMNs by MDSCs signifying a marked increase in efferocytosis (**Figure 5B**). Quantitative analysis was performed using flow cytometry which demonstrated a significant increase in uptake of apoptotic PMNs by WT M-MDSCs compared to *Mertk^-/-^* M-MDSCs (32.7±4.0 vs. 2.9±0.3; p<0.0001) (**Figure 5C-D**).

**Figure 5.**
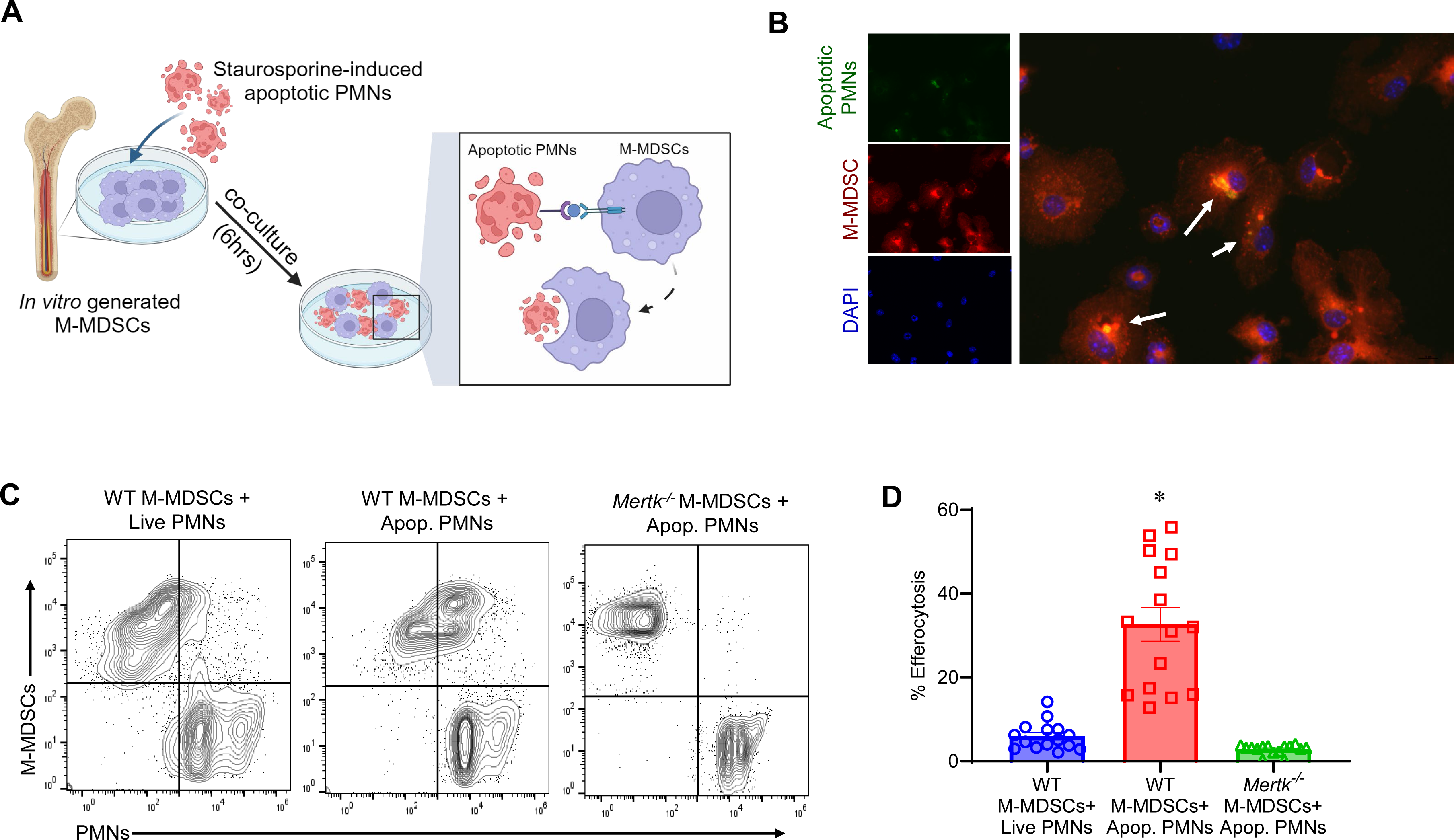
MerTK mediates M-MDSC-dependent efferocytosis *in vitro.* **(A)** Schematic depicting *in vitro* process of MerTK-mediated efferocytosis of apoptotic PMNs by M-MDSCs. **(B)** Representative immunofluorescence images demonstrating colocalization (yellow; white arrows) of M-MDSCs (red) engulfment of apoptotic PMNs (red). DAPI is shown in blue. **(C-D)**. Representative flow cytometry plots showing engulfment of apoptotic PMNs by WT or *Mertk^-/-^*-derived M-MDSCs. Quantification in lung tissue showed a significant increase in efferocytosis of apoptotic PMNs by WT mice-derived M-MDSCs but was absent in *Mertk^-/-^* mice-derived M-MDSCs. *p<0.0001 vs. all groups; n=15/group.

### Resolution of lung IRI in a murine orthotopic LTx model is mediated by M-MDSCs

To further validate the findings of the hilar ligation IRI model, we assessed the role of MerTK-dependent resolution of IRI in a clinically relevant murine BD orthotopic LTx model (**Figure 6A**). Following 72hrs of organ reperfusion there was a significant increase in M-MDSC infiltration compared to 24hrs of perfusion (8.8±0.9 vs. 3.1±0.2 %; p=0.02) (**Figure 6B-C**). This increase in M-MDSCs coincided with a significant decrease in albumin levels (0.4±0.04 vs. 1.1±0.09 ng/mL; p=0.006; **Figure 6D**) signifying decrease in lung edema, as well as proinflammatory cytokines (**Figure 6E-G**). Furthermore, there was enhanced efferocytosis during resolution evidenced by a significant decrease in PMN infiltration (216.7±34.5 vs. 765.9±49.7 PMNs/HPF; p=0.0014; **Figure 6H-I**), MPO expression (2301±929.5 vs. 8537±1131 pg/mL; p=0.014; **Figure 6J**), and sol-MER levels (257.7±40.4 vs. 831.3±65.17 pg/ml; p=0.003; **Figure 6K**) compared to 24hrs of reperfusion. Taken together, the results in both post-LTx models as well as *in vitro* studies outline the efferocytic role of M-MDSCs that immunomodulates the resolution of lung IRI (**Figure 6I**).

**Figure 6.**
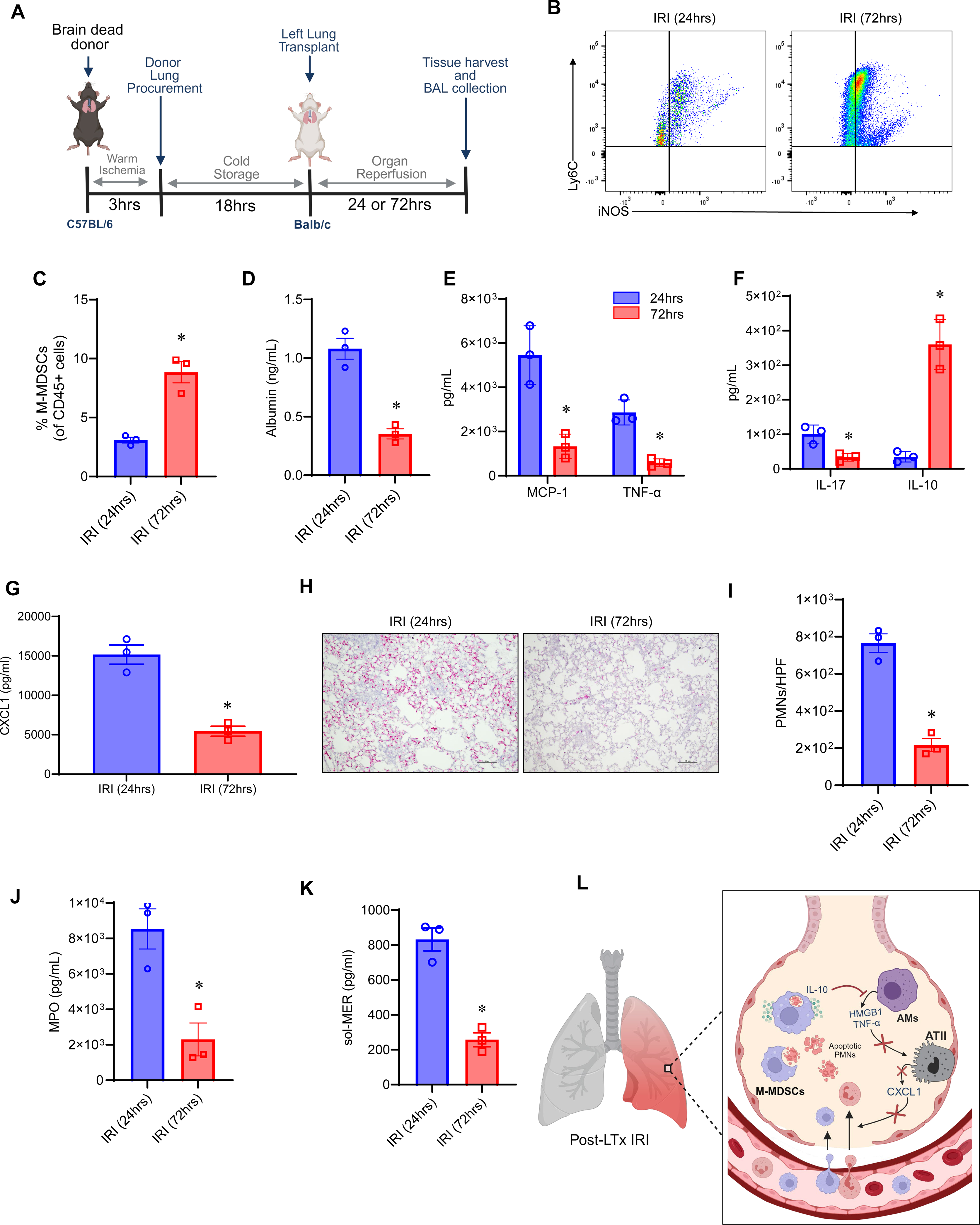
Resolution of IRI in an allogeneic orthotopic BD model of post-LTx IRI is associated with increase in M-MDSCs. **(A)** Schematic depicting brain dead (BD) model of LTx in C57BL/6 donors and WT (Balb/c) recipients. **(B-C)** Representative flow cytometry plots and quantification of M-MDSCs in lung tissue from LTx recipients. The percentage of M-MDSCs was significantly upregulated after 72hrs of reperfusion to 24hrs. *p=0.02 vs. 24hrs; *n*=3/group. **(D)** Albumin levels were significantly mitigated following 72hrs of reperfusion compared to 24hrs of reperfusion. *p=0.006; *n=*3/group. **(E-G)** Expression of pro-inflammatory cytokines was significantly mitigated and IL-10 expression was significantly increased after 72hrs compared to 24hrs of reperfusion. *p<0.04; *n*=3/group. **(H-I)** PMN infiltration was significantly abrogated following 72hrs of reperfusion compared to 24hrs *p=0.0014; *n*=3/group. **(J)** MPO expression was significantly mitigated following 72hrs of reperfusion. *p=0.014; *n*=3/group. **(K)** sol-MER levels were significantly decreased in LTx recipient mice following 72hrs of reperfusion compared to 24hrs. *p=0.003; *n*=3/group. **(L)** Schematic depicting the proposed immunosuppressive role of efferocytosis by M-MDSCs during the resolution of post-LTx IRI. M-MDSCs recognize, engulf apoptotic PMNs, and subsequently release anti-inflammatory IL-10 which then downregulates alveolar macrophage (AM) activation and inflammatory cytokine secretion. The cumulative effect of this signaling ameliorates alveolar type II (ATII)-mediated release of CXCL1, a potent neutrophil chemokine. Overall, upregulation of M-MDSC-mediated efferocytosis leads to inflammation-resolution via mitigating AM- and ATII cell-activation and downstream neutrophil transmigration during post-LTx IRI.

## DISCUSSION

The ongoing need to understand mechanisms of dysregulated resolution in PGD and subsequent CLAD development is evidenced by the 6.2 year median survival rate post-LTx (31, 32). The results reported herein describe a previously uncharacterized role of M-MDSCs in the resolution of lung IRI via efferocytosis of the apoptotic cell debris following post-LTx IRI. Since dysregulated clearance of dead cell debris can trigger uncontrolled inflammation and tissue injury, this process is crucial for effective resolution of lung injury and graft survival. We identified a significant role of the monocytic-cell population that can regulate efferocytosis in two murine LTx models as well as in human LTx samples. The importance and clinical relevance of these studies is indicated by the pivotal role of MerTK receptors in mediating efferocytosis and suggesting that prevention of MerTK cleavage on monocytic immune cell populations can alleviate post-LTx IRI.

The resolution of inflammation caused by IRI, or any sterile insult, is a highly coordinated and intricate process. Reparative mechanisms for homeostatic conditions include cessation of pro-inflammatory leukocyte infiltration, efferocytosis, and the production of pro-resolving molecules (i.e. anti-inflammatory cytokines, specialized pro-resolving lipid mediators, etc.), and the presence of pro-resolving/anti-inflammatory cell populations (6, 7). A particularly severe insult can disrupt one or more of these interdependent processes ultimately resulting in failed resolution. One notable pro-resolving cell population of the inflammatory response is the immunosuppressive M-MDSC subset that has pivotal immunoregulatory capabilities, and thus, can serve as a therapeutic modality in various disease conditions (33), However the contribution of M-MDSCs in the resolution of post-LTx IRI remains to be deciphered. While research surrounding the role of MDSCs in lung IRI is relatively unexplored, previous studies in other organ transplant models has investigated their relative contribution. A study of renal IRI found that depletion of both MDSC subsets via GR-1 antibody led to injury improvement and adoptive transfer of a pan population of MDSCs (including both G- and M-MDSC subsets) led to worsening of renal IRI (34). These results similarly support our own data where adoptive transfer of G-MDSCs provided no protection in lung IRI (data not shown). Instead of using strategies that deplete or adoptively transfer pan-MDSC population, we focused on specifically using the enriched MDSC subsets for our studies. This has allowed us to decipher the subset-specific contribution with specifically M-MDSCs contributing to inflammation-resolution.

Lung IRI is notably characterized by the infiltration of PMNs, which eventually undergoes cell death such as apoptosis or NETosis after the initial insult has subsided (27). The clearance, or efferocytosis, of these apoptotic cells is crucial for mitigating a feedforward loop of continuing inflammation and tissue injury (15). Though this process is under regulation by a variety of receptors, MerTK is an effective efferocytic receptor due to its high level of expression in multiple tissues of the body as well as on primary phagocytes like macrophages (35). MerTK expressed on the cell surface is subject to proteolytic cleavage by ADAM17, among other molecules, resulting in the generation of soluble-Mer (sol-Mer) (36). This not only decreases cell surface function of MerTK, but sol-Mer serves as a soluble ligand that further decreases ligand-dependent interactions and activation of MerTK receptor (28). Impaired MerTK function can lead to worsening of inflammation and dysregulated repair mechanisms in atherosclerosis, bacteria-induced lung injury, and myocardial IRI, whereas prevention of MerTK cleavage with genetic or pharmacological techniques has been demonstrated to enhance inflammation-resolution (37-39). Our results demonstrate the pivotal importance of preventing MerTK cleavage as a therapeutic strategy for mitigating lung injury and enhancing resolution.

Recent studies have elucidated the role of specialized proresolving lipid mediators (SPMs) such as RvD1 to prevent MerTK cleavage on macrophages and enhance efferocytosis (30, 40). It is important to note that various cell types are capable of performing MerTK-dependent or-independent efferocytosis, and the relative contribution of each cell is highly dependent on the injured microenvironment. Alveolar macrophages are the primary tissue resident professional phagocyte tasked with constant surveillance. Our previous studies have also demonstrated impaired efferocytosis of alveolar macrophages during peak inflammation (21). This is likely due to MerTK cleavage on alveolar macrophages during initial insult, whereas immune trafficking subsets, such as M-MDSCs, infiltrate the site of injury with functioning MerTK and thus effectively propagate efferocytosis. Additionally, its plausible M-MDSCs are acting in a multifaceted manner not only for efferocytosis, but through secretion of paracrine factors (i.e. cytokines, extracellular vesicles), which remains to be further elucidated. Beyond efferocytosis, maintaining or enhancing MerTK function during acute injury has the potential to enhance anti-inflammatory mediator production, anti-inflammatory macrophage function, and regulate macrophage lipid metabolism (30, 41- 43).

M-MDSCs are readily recruited to sites of inflammation through canonical monocyte trafficking pathways like the CCL2/CCR2 axis (44). Depending on the inflammatory microenvironment, these cells are capable of mediating immunosuppression through secretion of anti-inflammatory cytokines, recruitment of regulatory immune cells, exhaustion of pro-inflammatory cells through nutrient sequestering, and facilitating cell polarization to anti-inflammatory phenotypes (14). Previous studies have demonstrated the ability of MDSCs to prolong cardiac graft survival in a murine model, which was significantly reduced when MDSCs were depleted (45, 46), and M-MDSCs were shown to promote organ acceptance through recruitment of regulatory T cells in clinical kidney transplantation (47). Furthermore, studies in islet and heart transplantation have demonstrated an association between MerTK function and M-MDSC mediated transplant tolerance, namely through their ability to recruit regulatory T cells (48). Thus, our findings raise the interesting prospect of the role of MDSCs and MerTK for enhancing immunosuppression and alleviating post-LTx IRI.

There are limitations in this study that should be considered. The use of the hilar ligation model, which is self-resolving provides us with a high throughput way to assess the processes of resolution, but does not recapitulate the entire clinical process of LTx which includes cold preservation and donor-recipient characteristics that can further influence lung IRI. However, the use of allogeneic LTx model circumvents these concerns confirming the observed findings. Secondly, we did not use immunosuppressant therapy in the LTx model as our goal explored the inherent role of endogenous immune cell suppression by monocytic compartments without the influence of exogenous immunoregulation. Additionally, apart from MerTK, Tyro3 and Axl are a family of tyrosine kinase receptors, that may influence the process of efferocytosis and resolution (49). These receptors may act individually or concomitantly in a disease-dependent setting and should be deciphered in subsequent studies of allograft injury (50).

In summary, our findings establish a MerTK-mediated immunosuppressive mechanism that is novel to M-MDSCs in the resolution of lung IRI. This further adds to the growing body of evidence regarding the therapeutic potential of M-MDSCs, especially that can be enhanced *ex vivo*. Future clinical translational investigations should focus on enhancing the efferocytic ability of these cells through preservation of MerTK function.

## AUTHOR CONTRIBUTIONS

AKS designed the study. VL, DJMK, ZT, MWS, and AKS performed experiments. VL, DJMK, ZT, JAV, MWS, CA, LLM, PAE, GRU and AKS analyzed results. BS, MR and AE provided human BAL specimens. GC and JAV performed single cell sequencing analysis. VL and AKS prepared the manuscript with input from all authors.

## ACKNOWLEDGMENTS

The authors thank the ICBR core facility at UF (RRID:SCR_019119) and Ricardo Ungaro for assistance with flow cytometry experiments. Schematic figures were created using biorender.com.

