## Supplemental Methods for "MerTK-dependent efferocytosis by monocytic-MDSCs mediates resolution of post-lung transplant injury"

Victoria Leroy, Denny J.M. Kollareth, Zhenxiao Tu, Jeff Arni C. Valisno, Makena Woolet-Stockton, Biplab Saha, Amir M. Emtiazjoo, Mindaugas Rackauskas, Lyle L. Moldawer, Philip A. Efron, Guoshuai Cai, Carl Atkinson, Gilbert R. Upchurch, Jr., Ashish K. Sharma

**Supplementary Materials:**

1. **Supplementary Materials and Methods**
2. **Supplementary Figures**

**Supplementary Figure E1.** Dot plot of gene expression patterns utilized to identify Cluster 4 as M-MDSCs**.**

**Supplementary Figure E2**. mRNA expression of MerTK

**Supplementary Figure E3.** *In vitro* generation of M-MDSCs

**Supplementary Figure E4**. Resolution of lung IRI is impaired in *MerTK^-/-^*

**Supplementary Figure E5**. Pro-inflammatory cytokines are elevated in *MerTK^-/-^*

**Supplementary Figure E6**. Enhanced neutrophil infiltration and activation occurs during

failure of IRI resolution in *MerTK^-/-^* mice.

**Supplementary Figure E7**. MerTK-CR mice are protected following lung IRI

**Supplementary Figure E8**. MerTK-CR demonstrate protection in pro-inflammatory

cytokines

**Supplementary Figure E9.** MerTK-CR are protected against PMN infiltration and

activation

**Supplementary Figure E10**. Confirmation of MerTK mRNA expression.

3. **Supplementary Tables**

**Supplementary Table E1.** Efferocytosis-related genes

**Supplementary Table E2.** MerTK-efferocytosis related genes

**Supplementary Table E3.** Cluster 4 sequencing analysis

**Supplementary Table E4.** Murine flow cytometry antibodies

4. **References**

**SUPPLEMENTARY MATERIALS AND METHODS**

**Human LTx single-cell RNA sequencing analysis**

We analyzed scRNA-seq data of lungs explanted from four patients with chronic lung allograft dysfunction (CLAD) and three normal donors (DT). The original study published by Khatri et al., (GSE224210) (1) has annotated data of 40 cell populations in endothelial, epithelial, lymphoid, myeloid and mesenchymal lineages. In this study, we focused specifically on myeloid cell populations, which were filtered based on their original annotation, and reanalyzed cell clustering with the following steps: Data was analyzed using Seurat v5 after applying ‘nFeature_RNA > 250 & nFeature_RNA < 4000 & percent.mt < 15’ for quality control, 20,701 cells were included in further analysis (2). Data was normalized using the ‘LogNormalize’ method and CLAD and DT data were integrated using 30 identified anchors. Based on the top 20 principal components (PCs), the shared nearest neighbor (SNN) graph and Louvain algorithm were applied to identify 18 cell clusters, with a resolution parameter of 0.5. These cell clusters were visualized in a dimension-reduced space by the Uniform Manifold Approximation and Projection (UMAP) embeddings. We identified M-MDSC cells based on the expression of *HLA-DRA, ITGAM, CD33, CD14, FUT4, IL-10, VEGFA* (3, 4). Further, the CLAD data were compared to control data using Wilcoxon Rank Sum test and differentially expressed genes were detected by Log2FC>0.25, min.pct>0.01 and Bonferroni corrected p-value<0.05.

Sequencing analysis identified 12,061 genes, of which 1,940 were differentially expressed genes (DEGs) in CLAD vs. DT in M-MDSCs (cluster 4) (**Table E1**). We cross-referenced these genes with a curated list of efferocytosis-related genes (ERG) generated from a GeneCards® and PubMed® query (efferocytosis; **Table E2**). We further refined our analysis by isolating differentially expressed (DE-) ERGs with a secondary GeneCards query, specifically targeting MerTK-related efferocytosis genes (MerTK; efferocytosis; **Table E3**) and cross-referenced with this list. This dual-step approach ensured a more precise identification of genes related to both MerTK and efferocytosis within the dataset.

**Animal models of lung IRI**

An established murine *in vivo* murine left lung hilar ligation model was used with 8-12 week old male BALB/c (wild-type; WT), C/EBPβ^-/-^ (*cebpb*^-/-^), C57Bl/6, *Mertk^-/-^* (Jackson laboratory, Bar Harbor, ME), and MerTK-Cleavage Resistant (MerTK-CR; gift from Dr. Bishuang Cai, Icahn School of Medicine, Mt. Sinai, NY) mice as previously described (5, 6). Drinking water and standard chow diet were provided *ad libitum*.  All experiments were approved by and conducted in accordance to the Institutional Animal Care and Use Committee of the University of Florida (protocol#201810465). Briefly, mice were put under anesthesia with inhaled Pivetal® Isoflurane (Patterson Veterinary, Gainesville, FL). Following anesthesia, mice were intubated with PE-60 tubing and connected to a pressure-controlled ventilator (Harvard Apparatus Co, South Natick, MA) for Mechanical ventilation with room air performed at 150 strokes/min, 0.5 cc stroke volume, and peak inspiratory pressure <20 cm H2O. Mice underwent thoracotomy followed by left hilar occlusion for 1hr (ischemia) then reperfusion for 6- or 24hrs. Reperfusion was achieved by removing the hilar suture. Sham animals were subject to thoracotomy without hilar occlusion. *In separate groups,* mice were treated with 5.0x10^6^ M-MDSCs via intravenous (i.v.) injection 24hrs prior to IRI. To minimize ventilator-induced injury, mice were extubated and returned to their cage during ischemic and reperfusion periods. BAL fluid and left lung tissue was harvested after the end of respective reperfusion time points for further analyses.

Additionally, a murine orthotopic lung transplant was used with brain dead donors using C57BL/6 donors and Balb/c recipient mice, as previously described (7-9). C57BL/6 donor mice were anesthetized by intraperitoneal injection of ketamine and xylazine (0.01 mg/g) and ventilated with a mixture of isoflurane and oxygen. Donors received a tracheotomy followed by insertion of a ventilation cannula for mechanical ventilation of 120/min respiration rate and a tidal volume of 300-400µL. A paramedian borehole was used to insert a 4F balloon catheter. The balloon was slowly inflated over a 10-minute period resulting in irreversible ischemia via brain stem compression. Donor lungs were left untouched for a 3-hour period of “warm ischemia” which was followed by a flush with 2mL of 4°C Perfadex® through the main pulmonary artery. Donor lungs were then harvested and stored for 18hrs in Perfadex® at 4°C prior to transplantation into recipient mice. Left lungs were transplanted utilizing cuff techniques into Balb/c recipient mice. Following left lung transplants, mice were sacrificed after 24- or 72hrs and BAL as well as lung tissue was harvested for further analyses.

**Pulmonary function analysis**

Pulmonary function was analyzed via a buffer-perfused, isolated mouse lung system (Hugo Sachs Elektronik, March-Huggstetten, Germany), as previously described (10). After the reperfusion period, mice were anesthetized with ketamine and xylazine and a tracheostomy was performed. Ventilation of mice was provided with room air at 100 breaths/min at a tidal volume of 7μl/g body weight with a positive end expiratory pressure of 2 cm H_2_O. The lungs were perfused at a constant flow of 60μl/g body weight/min with Krebs-Henseleit buffer (Sigma-Aldrich, St. Louis, MO) containing 2% albumin 0.1% glucose and 0.3% HEPES (335–340 mOsmol/kg H_2_O). The lungs were equilibrated on the system for 5-minutes before hemodynamic and pulmonary parameters were recorded for an additional 5 minutes by the PULMODYN data acquisition system (Hugo Sachs Elektronik).

**Human bronchoalveolar lavage analysis**

Patients undergoing lung transplantation at the University of Florida were consented for Bronchoalveolar lavage (BAL) collection with Institutional Review Board approval (#IRB201900987). BAL collection was performed as part of routine surveillance bronchoscopy by sequentially instilling two 50 mL aliquots in the right middle lobe followed by aspiration immediately after instilling the second aliquot using syringe suctioning, as recommended by International Society for Heart and Lung Transplantation consensus statement for standardization of BAL in lung transplantation (11). Samples were taken immediately post-transplantation (day 0; donor) and on post-operative day 1 in sterile tubes. BAL was normalized for return volume collected, centrifuged at 500*g* for 5 min at 4°C and supernatants were utilized for sol-Mer analysis.

**Murine bronchoalveolar lavage analysis**

Following lung function analysis, right lungs were ligated and left lungs were lavaged with 0.3mL of cold PBS. The return fluid was centrifuged at 500g for 5min at 4°C. The supernatant was collected and stored at –80°C for future analysis.

**Cytokine and chemokine protein analysis**

A mouse-specific Bio-Plex Cytokine Assay panel (Bio-Rad, Hercules, CA) was used to quantify cytokine and chemokine protein content in murine BAL fluid, as per the manufacturer’s recommendations.

**Myeloperoxidase measurement**

Myeloperoxidase (MPO) was measured in murine BAL fluid using an MPO ELISA kit (R&D Systems, Minneapolis, MN), per the manufacturer’s instructions.

**Soluble MER Analysis**

Human and murine BAL was analyzed for quantification of soluble MER using a sol-MER kit (R&D Systems) per the manufacturer’s instructions.

**M-MDSC quantification via Flow Cytometry**

Left lungs of WT and *cebpb^-/-^* mice subject to sham or IRI surgeries were harvested and subject to enzymatic dissociation via Lung Dissociation Kit and subsequent mechanical dissociation via GentleMACS per manufacturer’s instructions (Miltenyi Biotec, Germany). The resulting cell suspension was filtered through a 70 µm cell strainer and then centrifuged at 300xg for 5 min at 4°C. Red blood cells were lysed with RBC Lysis Buffer (Roche, Millipore Sigma) by incubation for 2 min at 4°C. Following RBC lysis, cell suspensions were washed with PBS and centrifuged at 300xg for 5 min at 4°C. Cell pellets were resuspended in ice-cold PBS containing LIVE/DEAD™ Dye at 0.5µg per 1x10^6^ cells for 30 minutes at 4°C. Cells were then washed twice in ice cold PBS, and blocked with 0.5µg per 1x10^6^ cells mouse BD Fc Block™ (BD Biosciences, San Jose, California) diluted in eBioscience™ Flow Cytometry Staining Buffer (Invitrogen). After 5 minutes of blocking antibody, samples were stained with a cocktail of fluorophore conjugated-antibodies as described in **Table E4** for 30 min at 4°C. Cells were then fixed using eBioscience Intracellular Fixation and Permeabilization buffer set (eBioscience; Thermo Fisher Scientific) per manufacturer’s instructions. Cells were then subjected to intracellular staining of iNOS. Flow Cytometry was performed on a BD FACSymhony™A3 Cell Analyzer (BD Biosciences) and data was analyzed using FlowJo Software. M-MDSCs were identified as CD45^+^CD11b^+^CD11c^-^Ly6C^+^Ly6G^-^iNOS^+^ as previously described (4, 12) and quantified as a percentage of total CD45^+^ cells.

**Immunohistochemistry**

Following IRI, left lungs were immediately harvested and incubated in 10% neutral buffered formalin fixation (Sigma-Aldrich, St. Louis, MO) overnight. Tissues were embedded in paraffin and sectioned at 5µM. Neutrophil infiltration in lung tissue was performed by immunostaining lung sections with purified rat anti-mouse Ly-6G (Cat#551459, BD Biosciences, San Jose, CA). Alkaline phosphatase–conjugated anti-rat immunoglobulin G (Vector Laboratories, Burlingame, CA) was used as the secondary antibody, and signals were detected with Vector® Red substrate kit (Vector Laboratories). Sections were counterstained with Mayer’s Hematoxylin (Thermo Fisher Scientific, Waltham, MA). For each lung section, five random fields were captured at 20x. Neutrophil counts were acquired blindly via QuPath Software (13) and the counts were averaged per tissue sample.

***In vitro* generation of M-MDSCs**

M-MDSCs were generated using *in vitro* methods as previously described (14). Briefly, cells were isolated from bone marrow of WT, C57Bl/6, or *Mertk^-/-^* mice and cultured with granulocyte-macrophage stimulating factor (100ng or 2x10^3^ units per 3x10^6^ cells, Peprotech, Cranbury, NJ) in RPMI-1640 media containing 10% heat inactivated fetal bovine serum, 1% antibiotic-antimycotic, 1% L-glutamine, and 0.1% β-mercaptoethanol for 3 days. The resulting cell culture, consisting of M- and G-MDSCs, was further separated into Ly6G- and Ly6G+ subpopulations with Anti-Ly6G UltraPure Microbeads (Miltenyi Biotec) via AutoMACS magnetic based cell sorting. The Ly6G- population, representing M-MDSCs, was further cultured in the presence of LPS (0.1 ug/mL; Invitrogen) and IFN-γ (100 U/mL; Biolegend, San Diego, California) for 24hrs to induce immunosuppression. Immunosuppressive M-MDSCs were subject to a Dead Cell Removal kit via magnetic separation (Miltenyi Biotec) and live M-MDSCs were then subsequently used for *in vivo* and *in vitro* experiments.

***In vivo*** **efferocytosis**

Efferocytosis was evaluated separately in WT mice following sham, 6hr- or 24hr-IRI. In endogenous efferocytosis evaluation, staurosporine-induced (1µM, Cayman Chemicals, Ann Arbor, MI) apoptotic PMNs were labeled with pHrodo Red (Invitrogen) and administered at 1.0x10^6^ cells in 50µL of saline via intratracheal injection 1hr prior to ischemia. PMNs were acquired using a Neutrophil Isolation kit via AutoMACS (Miltenyi Biotec). Left lungs were harvested for efferocytosis analysis via flow cytometry. Endogenous M-MDSC efferocytosis was evaluated by identifying CD45^+^CD11b^+^CD11c^-^Ly6C^+^Ly6G^-^iNOS^+^PE^+^ populations representing M-MDSCs with engulfed PE-labeled apoptotic neutrophils. In exogenous efferocytosis evaluation, adoptively transferred M-MDSCs were labeled with CellTrace™ Violet (Invitrogen) and given via i.v. 24hrs prior to injury. Left lungs were harvested following IRI. Efferocytosis by exogenously administered M-MDSCs was evaluated by identifying populations double positive for CellTrace™ Violet and PE (pHrodo Red). Efferocytosis percentage was calculated by dividing double positive populations by total M-MDSCs. Flow cytometry was performed on a BD FACSymhony™A3 Cell Analyzer (BD Biosciences) and data was analyzed using FlowJo Software (BD Biosciences).

***In vitro* efferocytosis**

*In vitro* generated M-MDSCs from WT and *Mertk^-/-^* mice were co-cultured with live or staurosporine-induced apoptotic PMNs at a ratio of 1:3 for 6 hours and analyzed by flow cytometry. M-MDSCs were labeled with CellTrace™ Violet and PMNs were labeled with CellTrace™ Yellow Dye prior to co-culture. Efferocytosis was evaluated via flow cytometry for double positive populations. Also, *in vitro* generated M-MDSCs were stained with PHK26 Red Fluorescent Cell Linker (Sigma-Aldrich, St, Louis, MO) and plated at 0.2x10^6^ cells/well on collagen-gelatin coated culture chamber slides (Thermo Scientific, Waltham, MA) for 24 hours. PKH67 Green-labeled, staurosporine-induced apoptotic neutrophils were overlaid onto adherent M-MDSCs for 1hr. Chamber slides were washed with 1x PBS, fixed with 4% PFA, washed again with 1x PBS. Chamber slides were mounted to coverslips with Vectashield Vibrance mounting medium with DAPI (Vector Laboratories, Newark, California). Images were acquired at 40x using a Nikon Eclipse Ti-U microscope (Nikon, Tokyo, Japan).

**RNA Extraction, cDNA generation, and real-time quantitative PCR**

Total RNA was isolated from lung tissue and *in vitro* generated M-MDSCs using TRIzol according to the manufacturer’s instructions. The concentration of total RNA was determined using NanoDrop™ (Thermo Fisher Scientific; Grand Island, NY). iScript™ Reverse Transcription Supermix for RT-qPCR (Bio-Rad, Hercules, CA) was used to generate cDNA from 1000ng of total RAN. Subsequent detection and quantification of MerTK was performed using quantitative real-time PCR in a CFX96 Touch Real-Time PCR Detection System (Bio-Rad, Real-Time PCR Detection System (Bio-Rad,) using SsoAdvanced™ Universal SYBR Green Supermix according to manufacturer’s instructions. Assays were performed as follows: 1 cycle at 95°C for 2 min, then up to 40 cycles at 95°C for 05 sec and 63°C for 30 sec and then a melt curve was performed from 60°C to 95 °C (0.5 °C every 5 sec). For mRNA detection of MerTK, the following primers were used: MerTK (forward 5′-GCAAAAGTGACGTGTGGGCT-3′ and reverse 5′-GGGATCAGCACTCCAGCAAG -3′) and 18s (forward 5′-CGGCTACCACATCCAAGGAA -3′ and reverse 5′-AGCTGGAATTACCGCGGC-3′). All primers were synthesized by Millipore Sigma (St. Louis, MO). Results are reported as relative fold change and were calculated by utilizing the 2^(-ΔΔCt)^ method.

**Statistical analysis**

Statistical evaluation was performed with GraphPad Prism 10 software (GraphPad, La Jolla, CA). All values are presented as the mean ± standard error of the mean (SEM). One-way ANOVA followed by Tukey’s multiple comparison test was performed to compare differences between three or more groups, and T-test followed by Welch’s correction, Mann-Whitney test, or Wilcoxon rank sum test was used for pair-wise comparisons of groups. For data involving repeated measurements, two-way ANOVA with Geisser-Greenhouse correction, followed by Tukey’s multiple comparison test was used. A value of P< 0.05 was considered statistically significant.

**REFERENCES**

E1. Khatri A, Todd JL, Kelly FL, Nagler A, Ji Z, Jain V, Gregory SG, Weinhold KJ, Palmer SM. JAK-STAT activation contributes to cytotoxic T cell-mediated basal cell death in human chronic lung allograft dysfunction. *JCI Insight* 2023; 8.

E2. Hao Y, Stuart T, Kowalski MH, Choudhary S, Hoffman P, Hartman A, Srivastava A, Molla G, Madad S, Fernandez-Granda C, Satija R. Dictionary learning for integrative, multimodal and scalable single-cell analysis. *Nat Biotechnol* 2023.

E3. Song Q, Hawkins GA, Wudel L, Chou PC, Forbes E, Pullikuth AK, Liu L, Jin G, Craddock L, Topaloglu U, Kucera G, O'Neill S, Levine EA, Sun P, Watabe K, Lu Y, Alexander-Miller MA, Pasche B, Miller LD, Zhang W. Dissecting intratumoral myeloid cell plasticity by single cell RNA-seq. *Cancer Med* 2019; 8: 3072-3085.

E4. Bronte V, Brandau S, Chen S-H, Colombo MP, Frey AB, Greten TF, Mandruzzato S, Murray PJ, Ochoa A, Ostrand-Rosenberg S, Rodriguez PC, Sica A, Umansky V, Vonderheide RH, Gabrilovich DI. Recommendations for myeloid-derived suppressor cell nomenclature and characterization standards. *Nature Communications* 2016; 7: 12150.

E5. Cai J, Gehrau R, Tu Z, Leroy V, Su G, Shang J, Mas VR, Emtiazjoo A, Pelaez A, Atkinson C, Machuca T, Upchurch GR, Jr., Sharma AK. MicroRNA-206 antagomiR‒enriched extracellular vesicles attenuate lung ischemia‒reperfusion injury through CXCL1 regulation in alveolar epithelial cells. *J Heart Lung Transplant* 2020; 39: 1476-1490.

E6. Leroy V, Cai J, Tu Z, McQuiston A, Sharma S, Emtiazjoo A, Atkinson C, Upchurch GR, Jr., Sharma AK. Resolution of post-lung transplant ischemia-reperfusion injury is modulated via Resolvin D1-FPR2 and Maresin 1-LGR6 signaling. *J Heart Lung Transplant* 2023; 42: 562-574.

E7. Atkinson C, Floerchinger B, Qiao F, Casey S, Williamson T, Moseley E, Stoica S, Goddard M, Ge X, Tullius SG, Tomlinson S. Donor Brain Death Exacerbates Complement-Dependent Ischemia/Reperfusion Injury in Transplanted Hearts. *Circulation* 2013; 127: 1290-1299.

E8. Krupnick AS, Lin X, Li W, Okazaki M, Lai J, Sugimoto S, Richardson SB, Kornfeld CG, Garbow JR, Patterson GA, Gelman AE, Kreisel D. Orthotopic mouse lung transplantation as experimental methodology to study transplant and tumor biology. *Nat Protoc* 2009; 4: 86-93.

E9. Cheng Q, Patel K, Lei B, Rucker L, Allen DP, Zhu P, Vasu C, Martins PN, Goddard M, Nadig SN, Atkinson C. Donor pretreatment with nebulized complement C3a receptor antagonist mitigates brain-death induced immunological injury post-lung transplant. *Am J Transplant* 2018; 18: 2417-2428.

E10. Sharma AK, LaPar DJ, Zhao Y, Li L, Lau CL, Kron IL, Iwakura Y, Okusa MD, Laubach VE. Natural killer T cell-derived IL-17 mediates lung ischemia-reperfusion injury. *Am J Respir Crit Care Med* 2011; 183: 1539-1549.

E11. Martinu T, Koutsokera A, Benden C, Cantu E, Chambers D, Cypel M, Edelman J, Emtiazjoo A, Fisher AJ, Greenland JR, Hayes D, Jr., Hwang D, Keller BC, Lease ED, Perch M, Sato M, Todd JL, Verleden S, von der Thusen J, Weigt SS, Keshavjee S, bronchoalveolar lavage standardization w. International Society for Heart and Lung Transplantation consensus statement for the standardization of bronchoalveolar lavage in lung transplantation. *J Heart Lung Transplant* 2020.

E12. Eckert I, Ribechini E, Lutz MB. In Vitro Generation of Murine Myeloid-Derived Suppressor Cells, Analysis of Markers, Developmental Commitment, and Function. In: Brandau S, Dorhoi A, editors. Myeloid-Derived Suppressor Cells. New York, NY: Springer US; 2021. p. 99-114.

E13. Bankhead P, Loughrey MB, Fernández JA, Dombrowski Y, McArt DG, Dunne PD, McQuaid S, Gray RT, Murray LJ, Coleman HG, James JA, Salto-Tellez M, Hamilton PW. QuPath: Open source software for digital pathology image analysis. *Scientific Reports* 2017; 7: 16878.

E14. Eckert I, Ribechini E, Lutz MB. In Vitro Generation of Murine Myeloid-Derived Suppressor Cells, Analysis of Markers, Developmental Commitment, and Function. *Methods Mol Biol* 2021; 2236: 99-114.
