## Supplemental Figures for "MerTK-dependent efferocytosis by monocytic-MDSCs mediates resolution of post-lung transplant injury"

### Supplementary Figure E1

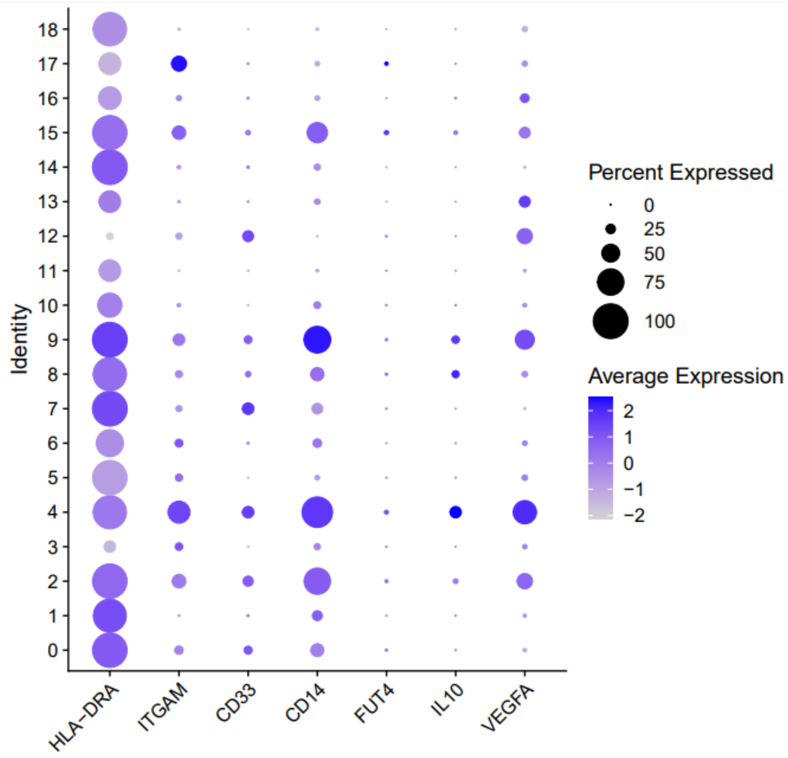


**Supplementary Figure E1.** Dot plot of gene expression patterns utilized to identify Cluster 4 as M-MDSCs based on expression of *HLA-DRA, ITGAM, CD33, CD14, FUT4, IL-10, VEGFA*.

### Supplementary Figure E2

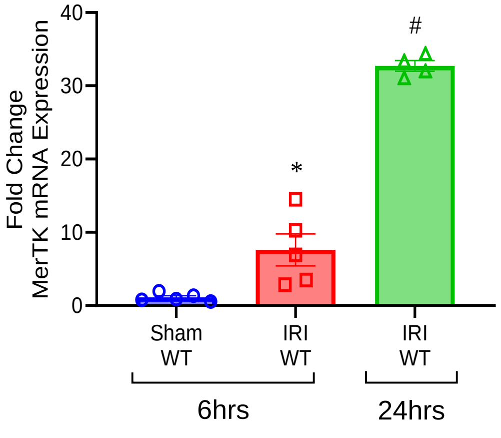


**Supplementary Figure E2**. mRNA expression of MerTK is significantly upregulated during inflammation-resolution phase of lung IRI. **(A)** A significant increase in MerTK expression was observed in WT mice during the resolution phase of lung IRI after 24hrs. *p<0.01 vs. sham; $^{\#}$p<0.001 vs. IRI (6hrs); $^{\delta}$p<0.001 vs. all groups. *n*=4-5/group.

### Supplementary Figure E3

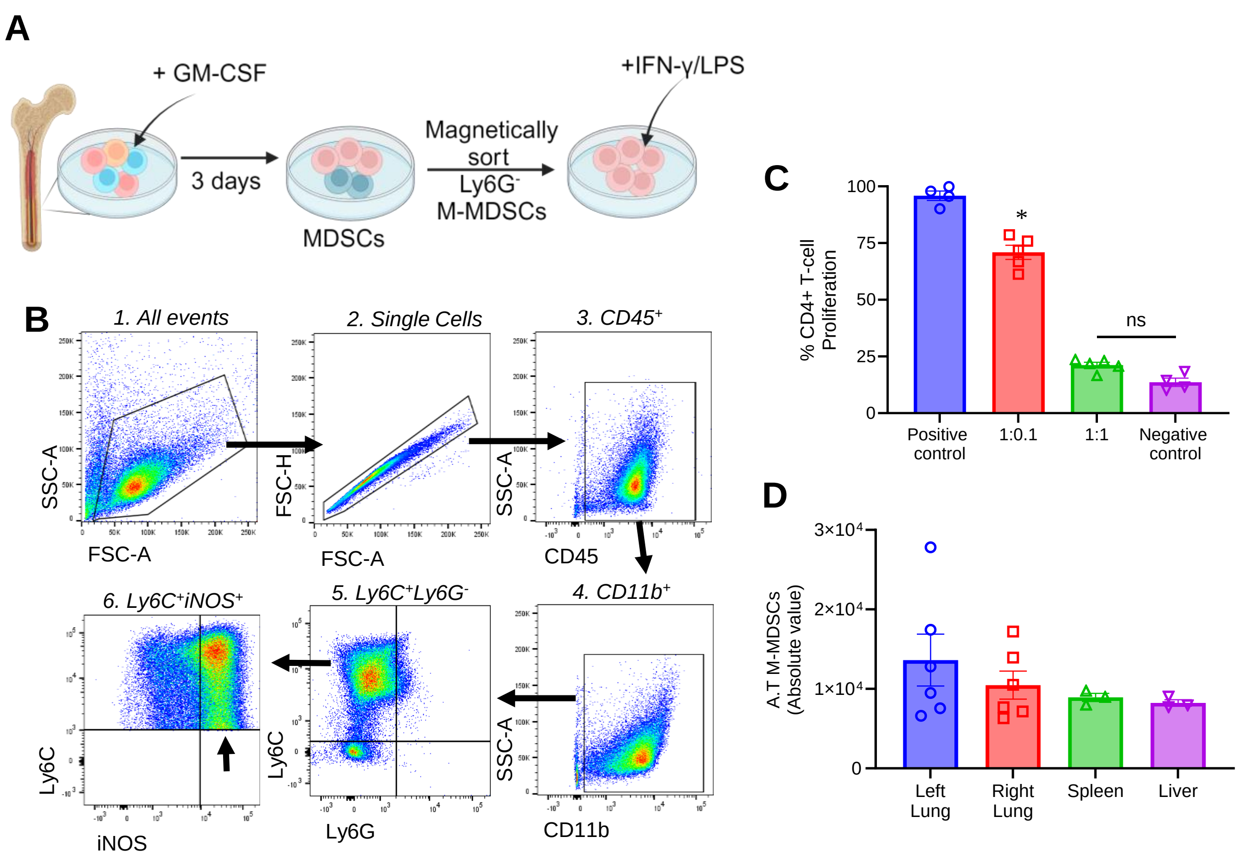


**Supplementary Figure E3**. *In vitro* generation of M-MDSCs. **(A)** Schematic depicting bone marrow harvest and *in vitro* culture to produce immunosuppressive M-MDSCs. **(B)** Representative gating strategy to confirm phenotype of M-MDSCs by $CD45^{+}CD11b^{+}Ly6G^{-}Ly6C^{+}iNOS^{+}$ cells. **(C)** Proliferation assays confirmed immunosuppressive ability of M-MDSCs to suppress $CD4^{+}$ T-cell proliferation in an M-MDSC dependent manner. Ratios are reported as T-cells:M-MDSCs. *p<0.001 vs. all other groups; ns, not significant; n=4-5/group. **(D)** Tracking of immunofluorescent labeled M-MDSCs after adoptive transfer confirm presence in the lung, spleen, and liver 24hrs after adoptive transfer administration. *n*=3-6 group.

### Supplementary Figure E4

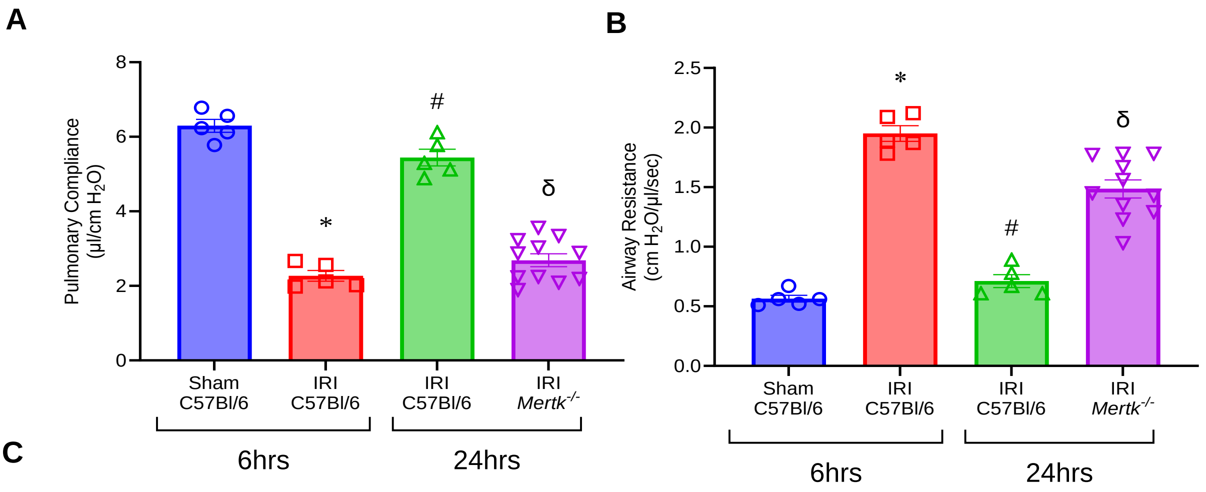


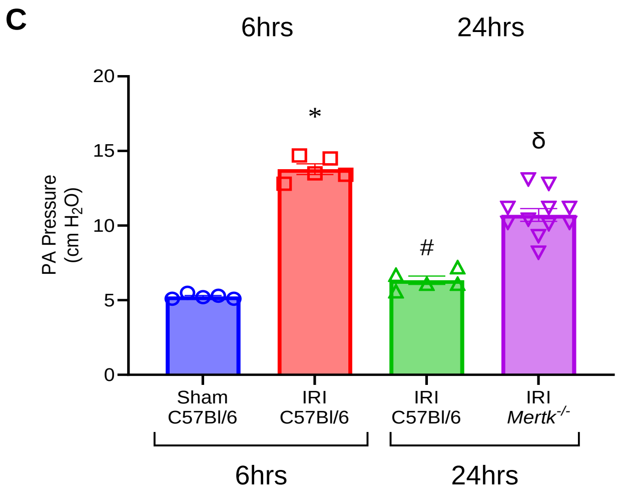


**Supplementary Figure E4**. Resolution of lung IRI is impaired in *Mertk*$^{-/-}$ mice*.* **(A-C)** Significant lung dysfunction was observed in *Mertk*$^{-/-}$ compared to C57Bl/6 mice after 24hrs. *p<0.0001 vs. sham C57Bl/6; $^{\#}$p<0.0001 vs C57Bl/6 IRI (6hrs); $^{\delta}$p<0.0001 vs. C57Bl/6 IRI (24hrs); *n*=5-7/group.

### Supplementary Figure E5

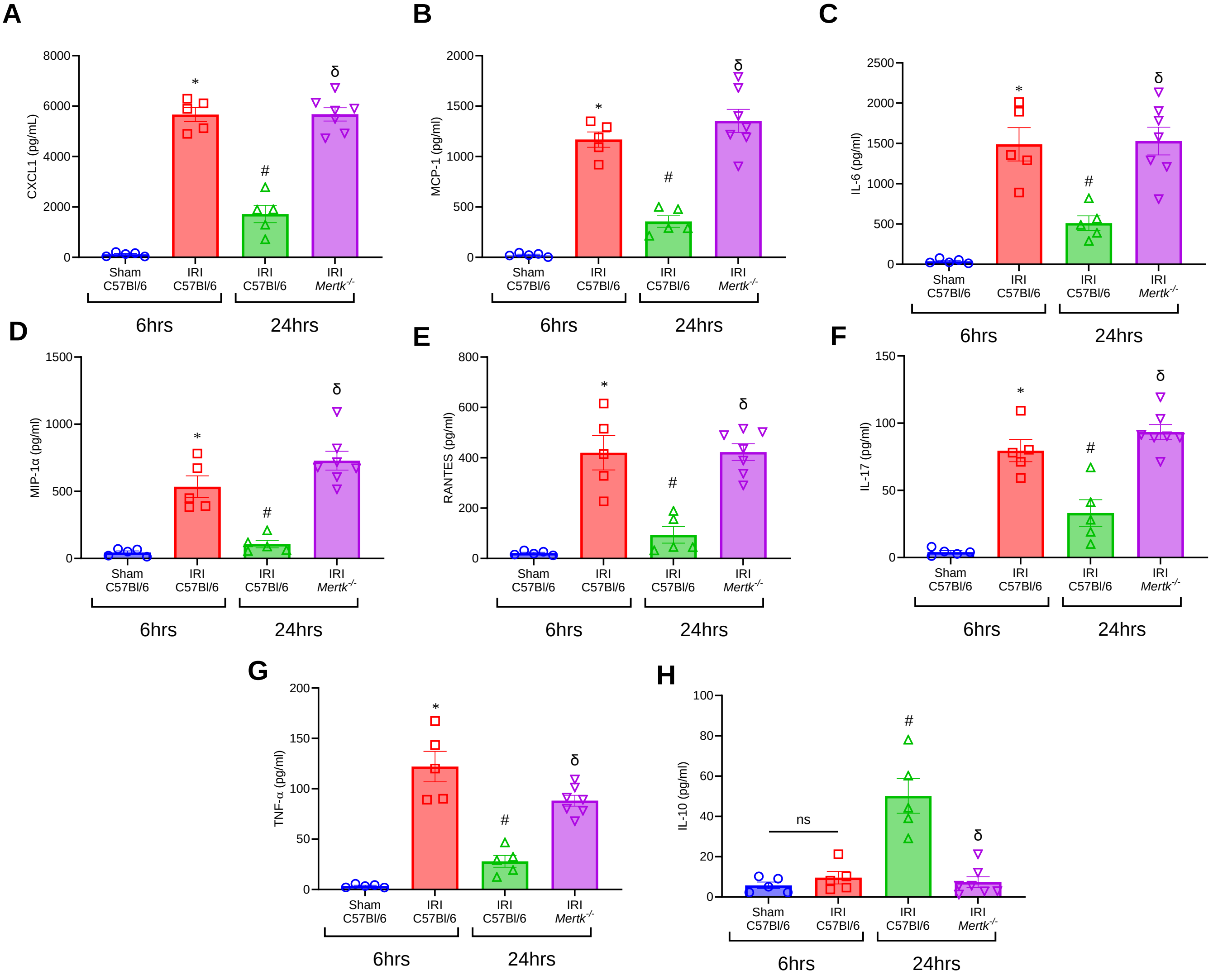


**Supplementary Figure E5**. Pro-inflammatory cytokines are elevated in *Mertk*$^{-/-}$. **(A-H)** Pro-inflammatory cytokine expressions in BAL were significantly increased and anti-inflammatory IL-10 expression was significantly decreased in *Mertk*$^{-/-}$ compared to C57Bl/6 mice after IRI (24hrs). *p<0.05 vs. sham C57Bl/6; $^{\#}$p<0.05 vs C57Bl/6 IRI (6hrs); $^{\delta}$p<0.05 vs. C57Bl/6 IRI (24hrs); *n*=5-11/group.

### Supplementary Figure E6

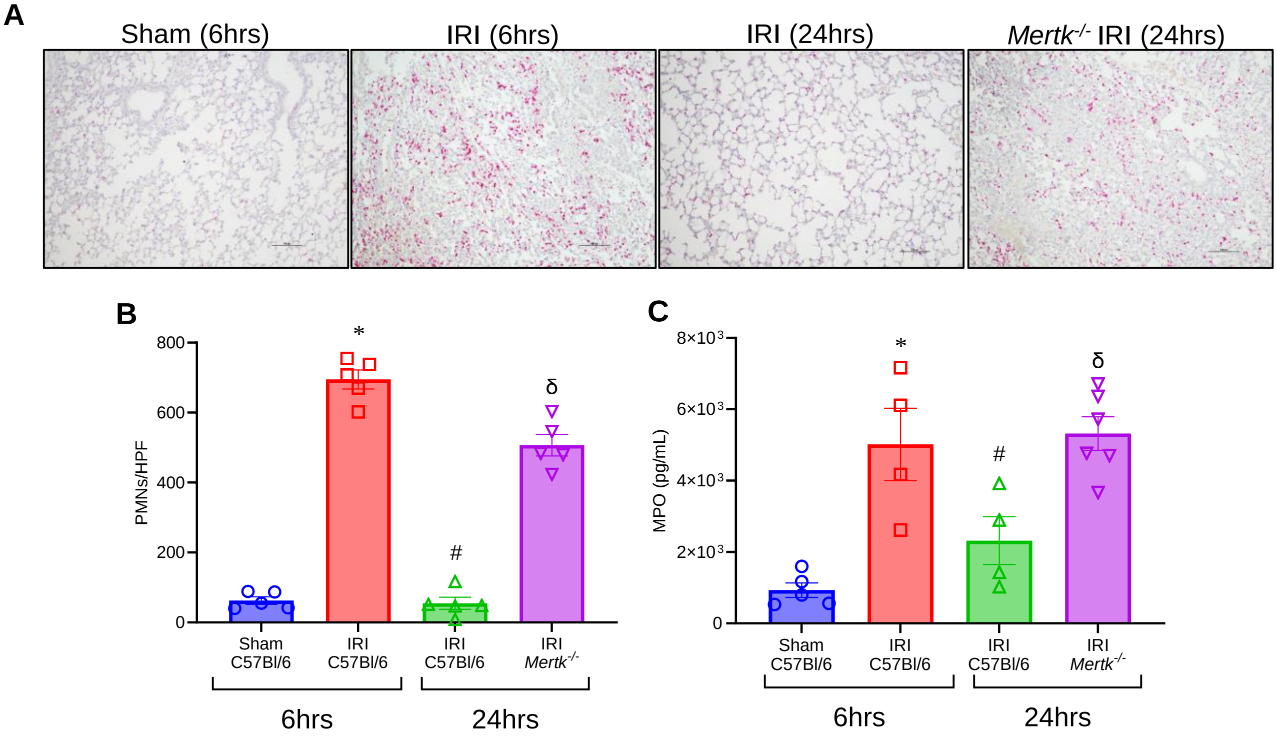


**Supplementary Figure E6**. Enhanced neutrophil infiltration and activation occurs during failure of IRI resolution in *Mertk*$^{-/-}$ mice. **(A-B)** Representative histological images and quantification of PMN infiltration demonstrate a significant increase in *Mertk*$^{-/-}$ mice compared to C57Bl/6 mice after 24hrs. *$p<0.0001$ vs. sham C57Bl/6; $^{\#}p<0.0001$ vs. C57Bl/6 IRI (6hrs); $^{\delta}p<0.0001$ vs. C57Bl/6 IRI (24hrs); *n*=5/group. **(C)** MPO levels remain significantly increased in *Mertk*$^{-/-}$ mice compared to C57Bl/6 mice after 24hrs. *$p<0.001$ vs. sham C57Bl/6; $^{\#}p=0.04$ vs C57Bl/6 IRI (6hrs); $^{\delta}p=0.01$ vs. C57Bl/6 IRI (24hrs); *n*=5/group.

**Supplementary Figure E7**

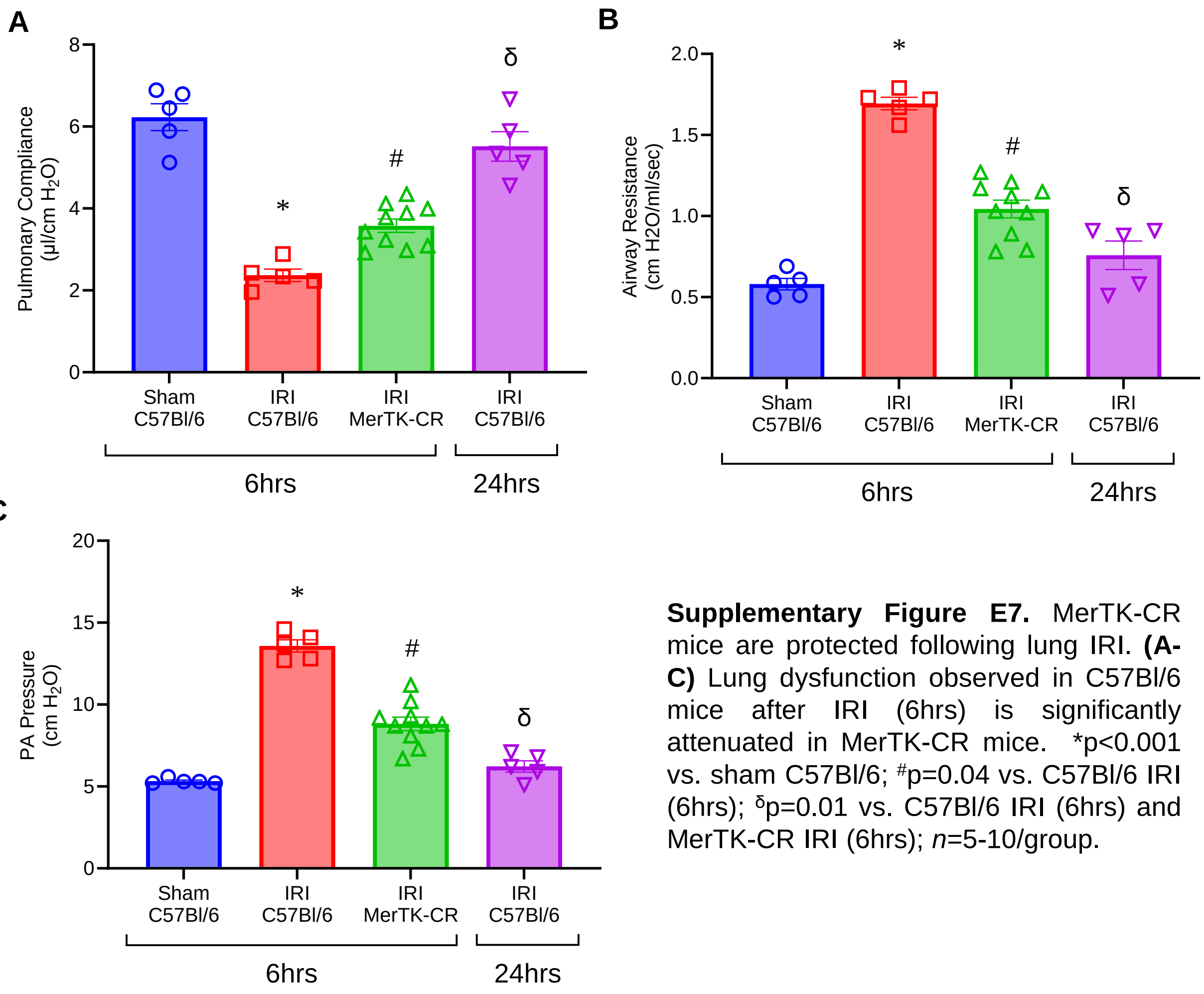


**Supplementary Figure E7.** MerTK-CR mice are protected following lung IRI. **(A-C)** Lung dysfunction observed in C57Bl/6 mice after IRI (6hrs) is significantly attenuated in MerTK-CR mice. *p<0.001 vs. sham C57Bl/6; $^{\#}$p=0.04 vs. C57Bl/6 IRI (6hrs); $^{\delta}$p=0.01 vs. C57Bl/6 IRI (6hrs) and MerTK-CR IRI (6hrs); *n*=5-10/group.

### Supplementary Figure E8

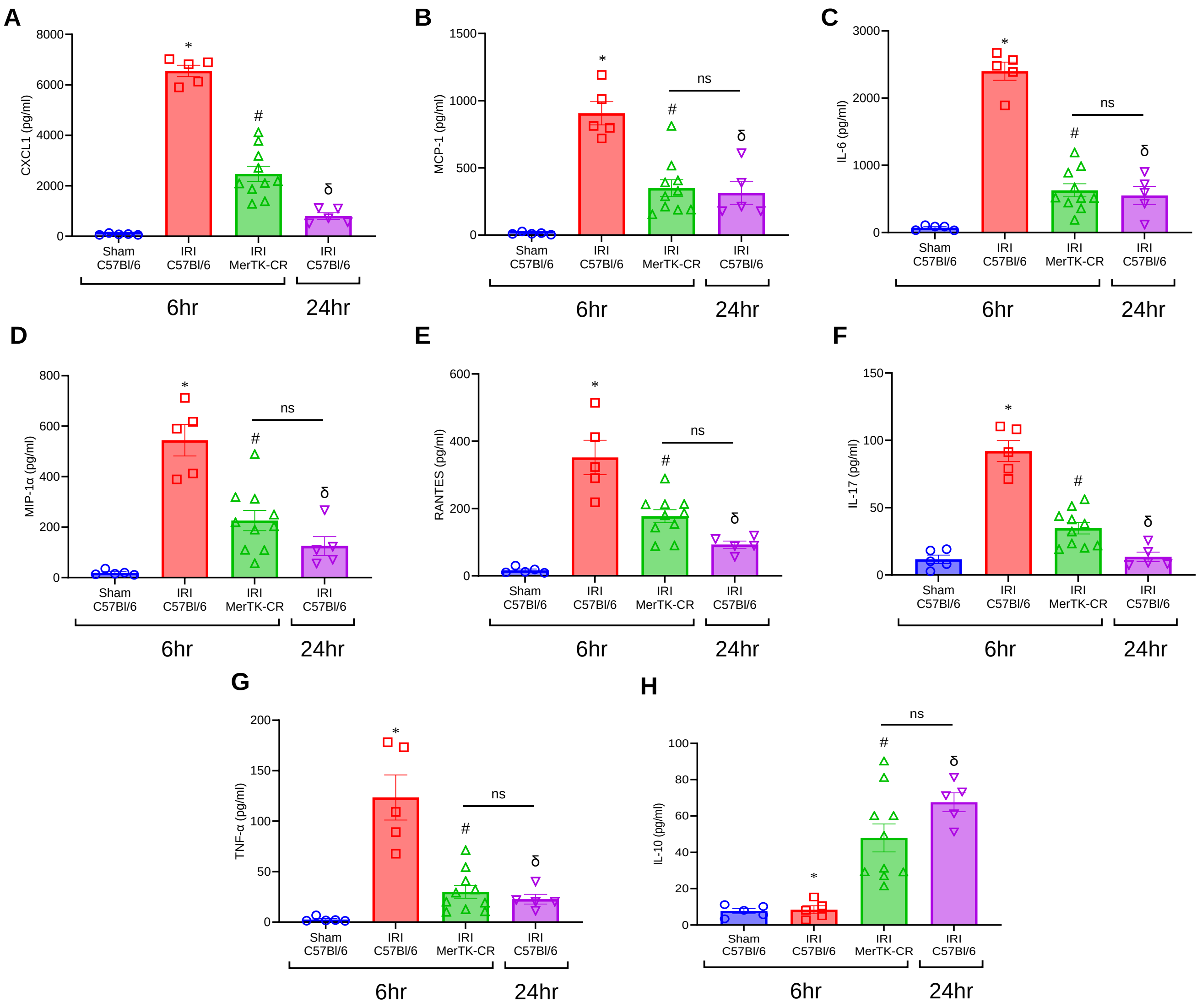


**Supplementary Figure E8**. MerTK-CR mice demonstrate protection in pro-inflammatory cytokines. **(A-H)** Similarly, MerTK-CR mice displayed significantly decreased pro-inflammatory cytokine and increased IL-10 expressions in BAL compared to C57Bl/6 mice following IRI (6hrs). ns, not significant; *p<0.05 vs. sham C57Bl/6; #p=0.05 vs. C57Bl/6 IRI (6hrs); δp=0.05 vs. C57Bl/6 IRI (6hrs) and MerTK-CR IRI (6hrs); *n*=5-10/group.

### Supplementary Figure E9

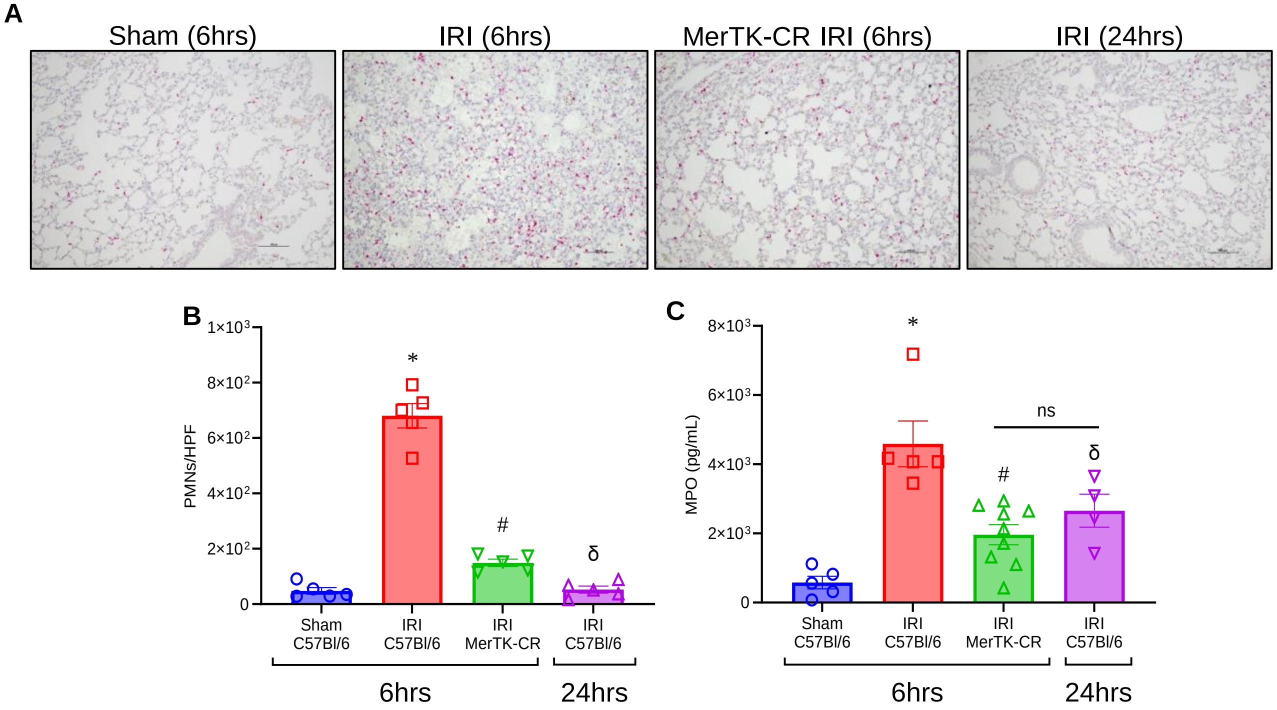


**Supplementary Figure E9**. MerTK-CR mice are protected against PMN infiltration and activation. **(A-B)** Histological staining for Ly6G and subsequent quantification revealed partial protection of MerTK-CR mice to PMN infiltration compared to C57Bl/6 mice following IRI (6hrs). *p<0.0001 vs. sham C57Bl/6; $^{\#}$p<0.0001 vs. C57Bl/6 IRI (6hrs); $^{\delta}$p=0.05 vs. C57Bl/6 IRI (6hrs) and MerTK-CR IRI (6hrs); *n*=5/group. **(C)** MPO levels were also significantly decreased in MerTK-CR mice after 6hrs compared to C57Bl/6. ns, not significant; *p<0.0001 vs. Sham C57Bl/6; $^{\#}$p=0.0006 vs C57Bl/6 IRI (6hrs); $^{\delta}$p=0.05 vs. C57Bl/6 IRI (6hrs); *n*=5/group.

**Supplementary Figure E10**

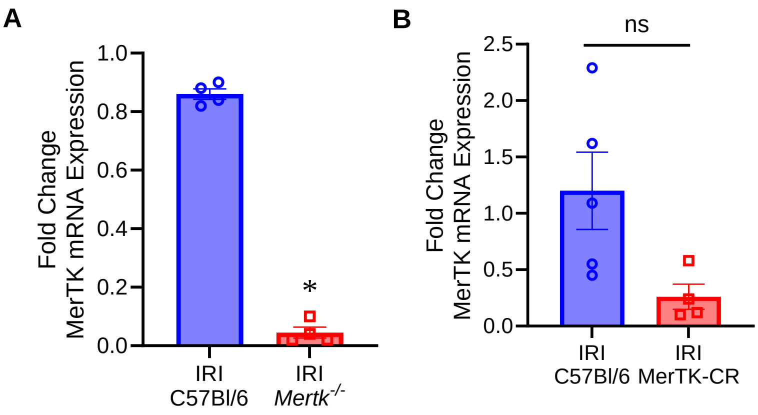


**Supplementary Figure E10**. Expression of MerTK in lung tissue of *Mertk*$^{-/-}$ and MerTK-CR mice. **(A)** mRNA expression showed *MerTK* was significantly down regulated following IRI (24hrs) in *Mertk*$^{-/-}$ mice as compared to C57Bl/6 mice. **(B)** MerTK mRNA expression in lung tissue after IRI (6hrs) remained the same in MerTK-CR mice as compared to C57Bl/6. ns, not significant; *$p<0.0001$ vs. C57Bl/6 IRI (24hrs); n=4-5/group.
